# Connecting natural and artificial neural networks in functional brain imaging using structured sparsity

**DOI:** 10.1101/390534

**Authors:** Christopher R. Cox, Timothy T. Rogers

**Author notes:** We have no conflicts of interest.

## Abstract

Artificial neural network models have long proven useful for understanding healthy, disordered, and developing cognition, but this work has often proceeded with little connection to functional brain imaging. We consider how analysis of functional brain imaging data is best approached if the representational assumptions embodied by neural networks are valid. Using a simple model to generate synthetic data, we show that four contemporary methods each have critical and complementary blind-spots for detecting distributed signal. The pattern suggests a new approach based on structured sparsity that, in simulation, retains the strengths of each method while avoiding its weaknesses. When applied to functional magnetic resonance imaging data the new approach reveals extensive distributed signal missed by the other methods, suggesting radically different conclusions about how brains encode cognitive information in the well-studied domain of visual face perception.

---

Scientists interested in understanding how minds emerge from brains have increasingly embraced a *network-based* view: cognition arises from the propagation of activity amongst distributed neural populations interacting via weighted connections in complex brain networks^1,2^. Information is encoded, not in the mean activation of local neural populations considered independently, but in the similarity structure of activity patterns distributed over many potentially distal populations^3,4^. This raises a statistical challenge for functional brain imaging, where various technologies yield thousands of neurophysiological measurements per second: cognitive structure must be encoded jointly over some *subset* of these measurements, but the number of possible subsets is prohibitively large. How can the theorist find those that encode structure of interest? We propose an answer motivated by neural network models of cognition^5^ and show that, when applied to data from functional magnetic resonance imaging (fMRI), the new approach can lead to dramatically different conclusions about the neural bases of cognition even in well-studied domains.

Neural network models (see SI-NN) propose that cognitive representations are patterns of activity distributed over neural populations or *units*, with each unit potentially contributing to many representations and each representation encoded over many units (Fig. 1A)^5,6^. The patterns arise as units communicate their activity through weighted *connections* that determine the effect of a sending unit on a receiving unit. Pools of similarly-connected units together function as a *representational ensemble* that encodes a particular kind of cognitive structure (phonological, semantic, visual, etc). Network topography is initially specified but learning shapes the connection weight values that generate patterns over ensembles. Cognitive processing arises from the propagation of activity through the network, typically from perceptual stimulation in a particular task and environmental context. Behaviors are the motor outputs eventually generated by such updating.

**Figure 1.**
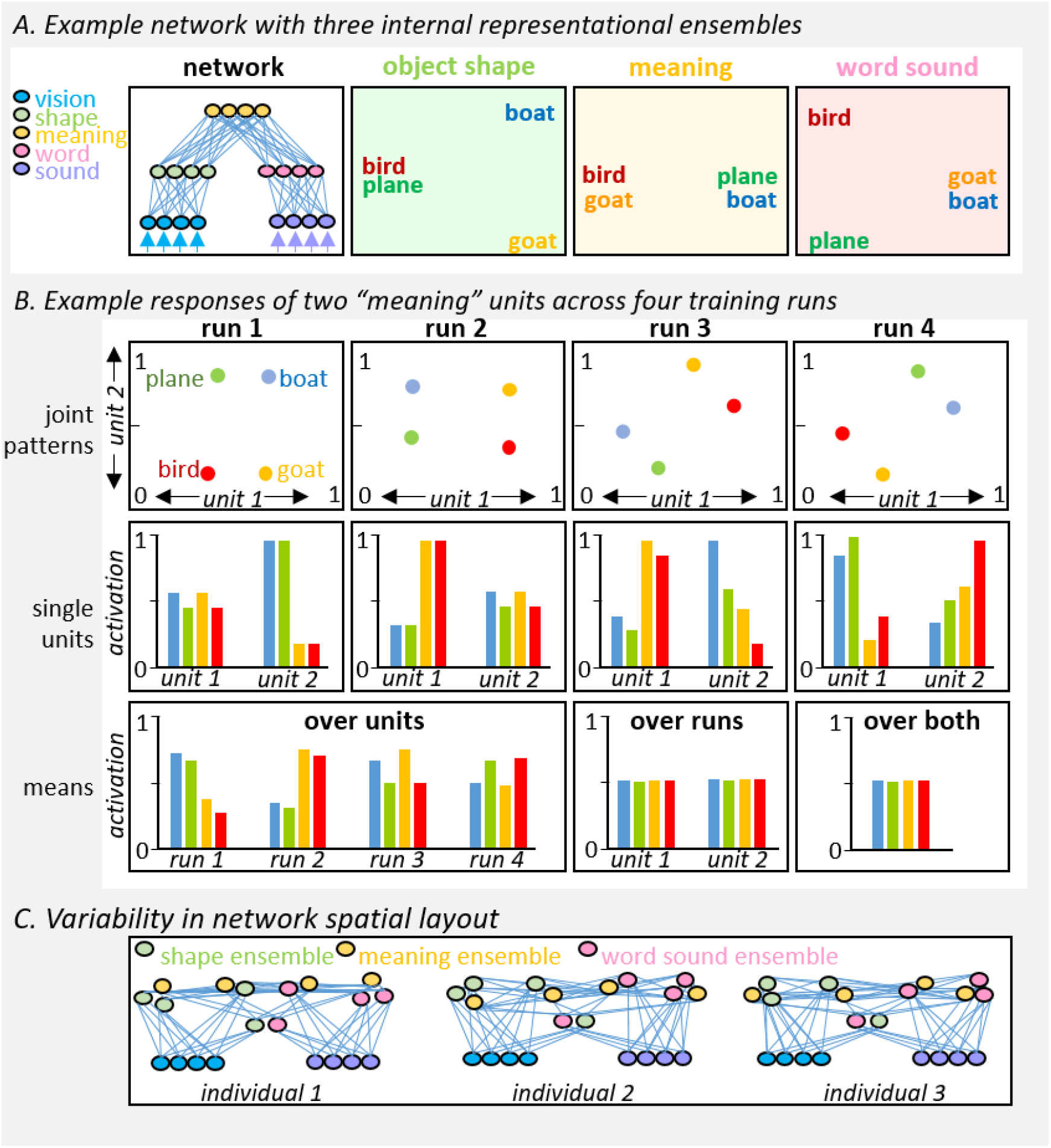
Challenges to functional brain imaging posed by network-based views of cognition. *A. Example network* that maps visual and auditory inputs to ensembles expressing similarity in shape, meaning, or word-sound, *B. Hypothetical contributions of two units* to the representation of meaning in such a network across four different training runs. Jointly the two units always encode the same distances amongst birds, goats, cars and boats (top row). Considered independently each unit appears to show a different response pattern both within and across runs (middle row). Consequently when the unit activations are spatially averaged within a run, or averaged across runs for each unit, their contributions to semantic structure are obscured (bottom row). *C. Variation in fine-grained connectivity*. All three depicted networks have the same connectivity as that shown in A and so will function the same way despite having a more complex spatial layout. The same coarse topography is expected across individuals but the finer-grained spatial layout—exactly where the green, yellow, and pink units lie within a given spatial volume—may vary, challenging approaches that average within region or across individuals.

Such models have long proven useful for understanding the neural bases of healthy^7^, disordered^8–11^, and developing^12,13^ cognition, but until recently^10,14–16^ this research advanced with little connection to brain imaging. The reason is simple—the models suggest that information can be encoded and distributed in ways that pose significant challenges to statistical analysis of brain imaging data (Fig. 1). Specifically:

1. *Unit activations may not be independently interpretable*. Because information resides in activation patterns computed across units in a representational ensemble, a single unit’s activation profile may mislead when considered in isolation (Fig 1B). Independent analysis of each population (e.g. single voxels) may therefore obscure the information encoded in distributed patterns.
2. *Neighboring units need not encode the same information in the same way*. Because representational structure inheres in pattern similarities over an ensemble, adjacent units in the ensemble need not respond to stimuli in similar ways—they may contribute to different components of the structure, or may encode a given component through either increased or decreased activation. Thus approaches that spatially average data within subject may destroy signal (Fig 1B bottom left).
3. *Single units can vary arbitrarily in their responses even when ensembles encode the same structure*. Many different weight configurations can compute the same input/output mapping for a given network. The configuration acquired through learning depends on factors that vary across individuals, such as the initial weight configuration or the idiosyncrasies of experience. Thus a particular unit in a given model can, across different training runs in the exact same environment, exhibit essentially arbitrary responses to inputs, even if the patterns of activation generated *across* unit subsets always express the same structure. Approaches that average voxel/region responses across subjects at a given anatomical location may therefore destroy signal (Fig 1B bottom middle and right).
4. *Representational ensembles need not be anatomically contiguous*. Depictions of network models portray representational ensembles as occupying a single contiguous “layer,” inviting a correspondence with cortical regions (Fig 1A). Yet units with similar connectivity will function as a representational ensemble—responding to the same inputs and contributing to the same outputs—even if situated in different cortical regions or intermingled with differently-connected units (Fig 1C). Thus approaches that analyze different anatomical regions independently can fail to uncover important signal.
5. *Fine connectivity varies across individuals*. While neural network models initially specify coarse connectivity, the values ultimately taken by individual weights are seeded randomly then tuned by learning, similar to real brains where region-to-region connectivity is initially specified but finer patterns are more variable and plastic. Consequently precise unit-to-unit alignment may be impossible even if the neural populations contributing to some representation reside in roughly similar locations across individuals (Fig 1C)—in which case cross-subject averaging of neural responses can destroy signal even with sophisticated alignment techniques.

To understand whether and how contemporary methods overcome these challenges, we first describe a simulated functional imaging study using a simple neural network to generate model data, which we then analyze with four different statistical methods. Each discerns some model signal but also has critical blind spots. The contrasting patterns suggest a new approach based on *structured sparsity*^17^ that preserves the strengths of other methods while avoiding their weaknesses. We develop this approach, then compare it to the others when applied to real fMRI data to discover neural signal that discriminates visually-presented faces from other visual stimuli. Each method again yields different results, but the new approach resolves seeming discrepancies amongst the others while simultaneously suggesting radically different conclusions about how neural systems encode representations in this well-studied domain.

## Results

### Simulation study

The model (Fig 2A) is an auto-encoder: it learns to reproduce input patterns across output units. The patterns come from two domains, A and B, corresponding to some cognitive distinction of interest (e.g. faces vs places, animals vs artifacts, etc). Each domain exemplar generates a unique pattern of input/output activations. *Systematic input/output (SIO) units* each independently adopt a noisy but consistent code, with some slightly more active on average for A items and others for B items. *Arbitrary input-output (AIO) units* have equal probability of activation regardless of domain. Activation propagates from input to output via two hidden layers. The *systematic hidden (SH) units* connect only systematic I/O units, while the *arbitrary hidden (AH) units* receive connections from arbitrary inputs and send connections to both systematic and arbitrary outputs. This architecture promotes a division of labor across the hidden units: SH units encode distributed representations that strongly express domain structure, while AH units encode the idiosyncratic differences existing amongst domain exemplars. The model also contains *irrelevant units* unconnected to the network and taking on low random values.

**Figure 2.**
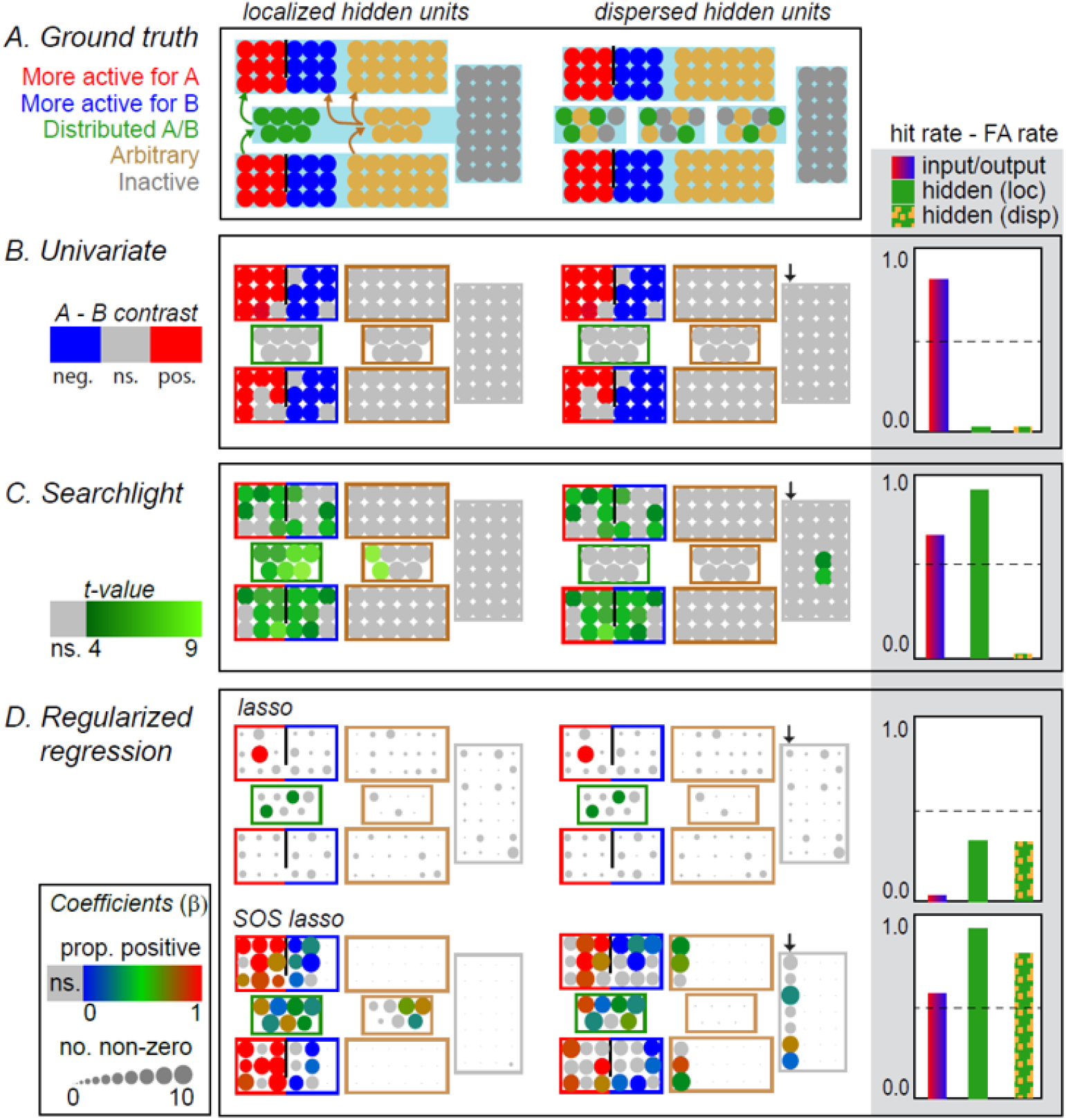
Simulation results. *A. Model architecture*. Circle colors indicate the unit type. Arrows in the left panel indicate connectivity, which is the same in localized and dispersed layouts. Blue shading indicates units treated as anatomically contiguous. *B-D. Results for each statistical method*. Colored circles indicate units identified as significantly contributing to the A/B distinction with p < 0.002 uncorrected. For regularized regression, circle size indicates the number of times the unit received a non-zero coefficient across 10 model runs. For units identified as important, the direction of the effect is plotted as a heat map with hot units activating more for A items, cool units activating more for B items, and green units showing uncertain or mixed direction of effect. Layer border colors indicate the unit type according to the legend in panel A. Hidden units in the localized and dispersed cases have been laid out in the same way to facilitate comparison of results. In each plot, a successful method will identify all units in the three leftmost boxes and no units in the remaining boxes. For the dispersed layout, the arrow indicates the column of irrelevant units that were intermixed with arbitrary and systematic hidden units. Barplots show how accurately each method discriminates signal-carrying from arbitrary and irrelevant units, measured as the hit rate minus the false alarm rate, and computed separately for input/output units (red/blue bar), hidden units in the localized layout (green bar), and hidden units in the dispersed layout (green/yellow bar).

Blue shading in Figure 2A shows units treated as anatomically contiguous. I/O units of a given type (A, B, arbitrary) were always anatomically grouped in the same way across model individuals within a single contiguous region. For the hidden and irrelevant units we considered two spatial layouts. When *localized*, units of a given type (SH, AH, irrelevant) were anatomically grouped in the same way across model individuals within a single contiguous region, just like I/O units. When *dispersed*, the model retained the same connectivity but the hidden units were spatially arranged in four anatomically distal “regions”—three containing a mix of SH, AH, and irrelevant units and the fourth containing only irrelevant units—to capture the possibility that distal units can function together as a representational ensemble. Within each region, unit locations were shuffled for each model subject, capturing the possibility that fine-grained localization can vary across individuals. These different configurations did not affect model training or behavior, but represent different assumptions about the spatial locations of the signals measured in a simulated MRI study.

Simulated data were generated by training a model on 72 item, then computing the response of each unit to each input pattern and perturbing this with noise sampled independently for each unit. Each unit activation was taken as a model analog of the peak BOLD signal generated by a single stimulus at a single voxel in a single individual. To simulate 10 individuals in a brain imaging study we trained and tested the model 10 times with different initial weights. We then applied four different statistical methods to find units that differentiate A from B items: univariate contrast (UC), searchlight multivariate pattern classification (sMVPC), and whole-brain logistic regression regularized with the L2 (ridge) or L1 (LASSO) norm (see SI-Approaches). We measured how well each method discriminates informative from non-informative units amongst all I/O units and amongst all hidden/irrelevant units. We also considered how well each method uncovers the domain code: which units are more active for A, which for B, and which adopt a heterogeneous code. Figure 2 shows the results.

*Univariate contrast* spatially smooths data, then identifies voxels whose mean activation across individuals at a given spatial location reliably differs for A vs B items.^18,19^ This approach identified the SIO units and accurately specified their coding direction, but failed to identify the SH units in either spatial layout.

*Searchlight MVPC* generates *information maps* by evaluating pattern classifiers in their ability to discriminate A from B items from the activation patterns generated by each item across voxels^20^. The approach avoids model over-fitting by applying the classifier only to voxels within a *searchlight* of fixed anatomical radius centered on a particular voxel. A separate classifier is evaluated for each voxel in each subject without spatial smoothing or cross-subject averaging. Cross-validation accuracy is stored at the voxel location in each subject; cross-subject univariate tests then identify anatomical clusters where accuracy is reliably higher than chance. The resulting maps indicate regions where the categorical distinction of interest is *locally decodable* on average across subjects within the radius of the searchlight. Figure 2C shows model results for the best-performing searchlight size (radius 7) which excelled at discovering localized signal in both I/O and hidden units. Smaller searchlights successfully identified SH but not SIO units while larger searchlights showed the reverse pattern (see SI-Approaches). Regardless of size, the approach failed to discover signal-carrying hidden units when these were spatially dispersed. Also, because information maps only indicate classifier accuracy, the approach did not reveal the different category codes employed by I/O and hidden units.

*Regularized whole-brain MVPC* fits and evaluates a single classifier per subject using all voxels and avoids over-fitting through model regularization^21^ –that is, by finding classifier coefficients that jointly minimize both prediction error and some additional penalty. We used logistic regression as the classifier and considered two common regularizers: the *ridge* penalty, which increases with the sum of squared classifier coefficients, and the *LASSO* penalty, which increases with the sum of the absolute value of classifier coefficients. In both cases a classifier was fit to minimize:

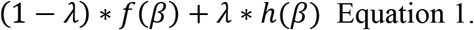

…where β is the vector of classifier coefficients (one element for each unit), *f*(*β*) is the classifier logistic loss, *h*(*β*) is the regularization penalty, and *λ* ∈ [0,1] is a free parameter tuned via cross-validation that determines the relative weighting of the loss vs the regularization penalty.

The ridge classifier showed a mean hold-out accuracy of 60% correct, reliably above chance (p < 0.0001), but placed non-zero weights on all units making it difficult to discriminate signal-carrying from arbitrary/irrelevant units (see SI-Approaches). The LASSO classifier showed equivalently good hold-out accuracy (60%) with a much sparser solution (Fig. 2D). Only 2 of the 7 SH units and only one of the I/O units were selected more often than expected by chance given the base probability of selection in a permutation test, and no false alarms were observed. Thus LASSO showed high precision—all selected units carried signal—with a low hit rate.

*Regularization with structured sparsity*. These results highlight complementary strengths and weaknesses across methods. Spatial averaging allows UC to detect the noisy signal in SIO units and also reveals the code direction, but only succeeds when units employing the same code are consistently localized within a region across subjects—because SH units employ a heterogeneous code, they are always missed. When sized appropriately, searchlight can discover both kinds of localized signal but misses anatomically dispersed signal and obscures the code direction. Regularized whole-brain MVPC can identify spatially dispersed and heterogeneous signal, but ridge regularization selects everything without discrimination while LASSO forces a very sparse solution that identifies only a small proportion of signal-carrying units.

Structured sparsity^22–24^ provides an avenue for preserving the strengths of each method while avoiding its weaknesses. A single classifier is fit using all data from all subjects, but the fit is regularized with a penalty that encourages desired sparsity patterns amongst the classifier coefficients. Specifically, the solution should reflect the characteristics of neural signal that neural network models suggest and that other methods individually exploit. It should (1) clearly delineate selected and unselected voxels, (2) allow heterogeneous codes among neighboring units within and across individuals, (3) reveal code direction where this is consistent, (4) identify distal units that jointly express representational structure, (5) capitalize on shared location across subjects where this exists but also (6) tolerate individual variation in signal location.

The *sparse-overlapping-sets* (SOS) LASSO is one such function^25,26^ (see SI-SOS). Neighboring voxels within a specified radius are grouped into *sets*, similar to searchlights. Each voxel belongs to several sets and sets overlap in the voxels they contain. Sets then contribute to the regularization cost of a single classifier as follows:

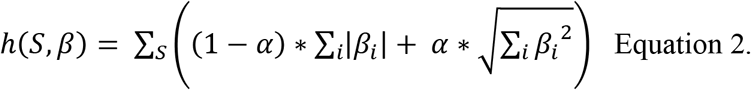

…where *S* defines the grouping of voxels into sets, *i* indexes model coefficients within a set, and *α* ∈ [0,1] is a free parameter. The total cost is a sum over sets. The cost for each set is the proportional weighted sum of two terms: the standard L1 *sparsity* penalty, and a *grouping* penalty formulated as the root of the sum of squared coefficients. Because this root is taken over units within a set, the grouping penalty is smaller when non-zero coefficients occupy the same set than when they occupy different sets (see SI-SOS). Thus SOS LASSO encourages sparse solutions where selected voxels occupy a small number of sets. The free parameter α controls the relative weighting of the grouping vs sparsity penalties within set—when α=0 the penalty reduces to LASSO. This function replaces *h*(*β*) in Equation 1, so the full optimization jointly minimizes model error and regularization cost with a second free parameter λ controlling their respective weighting.

SOS LASSO fits a single classifier to all data from all subjects simultaneously. Voxels from different subjects are projected into a common reference space without interpolation and with no cross-subject averaging. Sets are defined for grid points in the space and each encompasses all voxels within and across subjects that fall within a specified radius of a grid point. The optimization is convex and returns a unique solution for a given pair of hyperparameters (α and λ) that are tuned via cross-validation on model hold-out error. The result is a classifier coefficient for each voxel in each subject, with the coefficients tending to occupy a small number of sets, and thus to lie in roughly similar anatomical locations across subjects.

Like LASSO, SOS LASSO “sees” all voxels at once and so can exploit structure coded across distal regions, while also clearly delineating selected voxels by forcing many coefficients to zero. Like searchlight, the code direction can be heterogeneous within region and across subjects, but like LASSO each voxel in each subject gets a unique coefficient whose sign indicates the code direction. SOS LASSO promotes a similar anatomical distribution of voxels across subjects, but the pressure to be similar is balanced with pressure to reduce model prediction error and achieve structured sparsity within each subject. Since the strength of the grouping penalty is tuned by data, the approach can exploit localization across subjects where this exists but can also accommodate cross-subject variability in location.

The last row of Fig 2D shows the SOS-LASSO solution applied to model data. The classifier achieved better hold-out accuracy than LASSO or ridge regression (70% correct) while discovering more than half the SIO units and almost all the SH units in both localized and dispersed layouts. The only false alarms were observed amongst arbitrary or irrelevant units that were anatomically interspersed with or neighboring the signal-carrying units. The signs of the coefficients revealed the consistency and direction of the domain code among identified SIO units as well as the heterogeneity of the code in SH units. Thus in simulation, SOS-LASSO captures the strengths of each method while largely avoiding their limitations.

### Functional imaging study

The simulations suggest that several contemporary statistical methods for functional imaging have blind spots, raising the possibility that prior work may have missed important signal even in well-studied domains where multiple different approaches have been applied. Study 2 assesses this possibility in the domain of face representation. We applied univariate contrast, searchlight MVPC, LASSO, and SOS-LASSO to fMRI data collected in a previous unrelated study^27^ in which 10 participants judged the pleasantness of images depicting people (30 items), scenes (30 items), or objects (30 items) while their brains were scanned with fMRI. The study employed a slow event-related design that allowed estimation of the peak BOLD response to each item at each voxel without time-series deconvolution. We applied each method to these data to find voxels that reliably discriminate face (people) from non-face (place and object) stimuli.

Figure 3 shows the neural systems typically thought to support face and place perception (downloaded from http://web.mit.edu/bcs/nklab/GSS.shtml) along with the results for the current data from each method. Whole-brain univariate contrast revealed significantly reduced mean activation for faces in bilateral parahippocampal regions (blue areas p < 0.05 with cluster correction), while an ROI analysis found reliably elevated activation for faces in the right “fusiform face area” (FFA) only (warm colors, p < 0.05 for mean response across ROI voxels).

**FIGURE 3.**
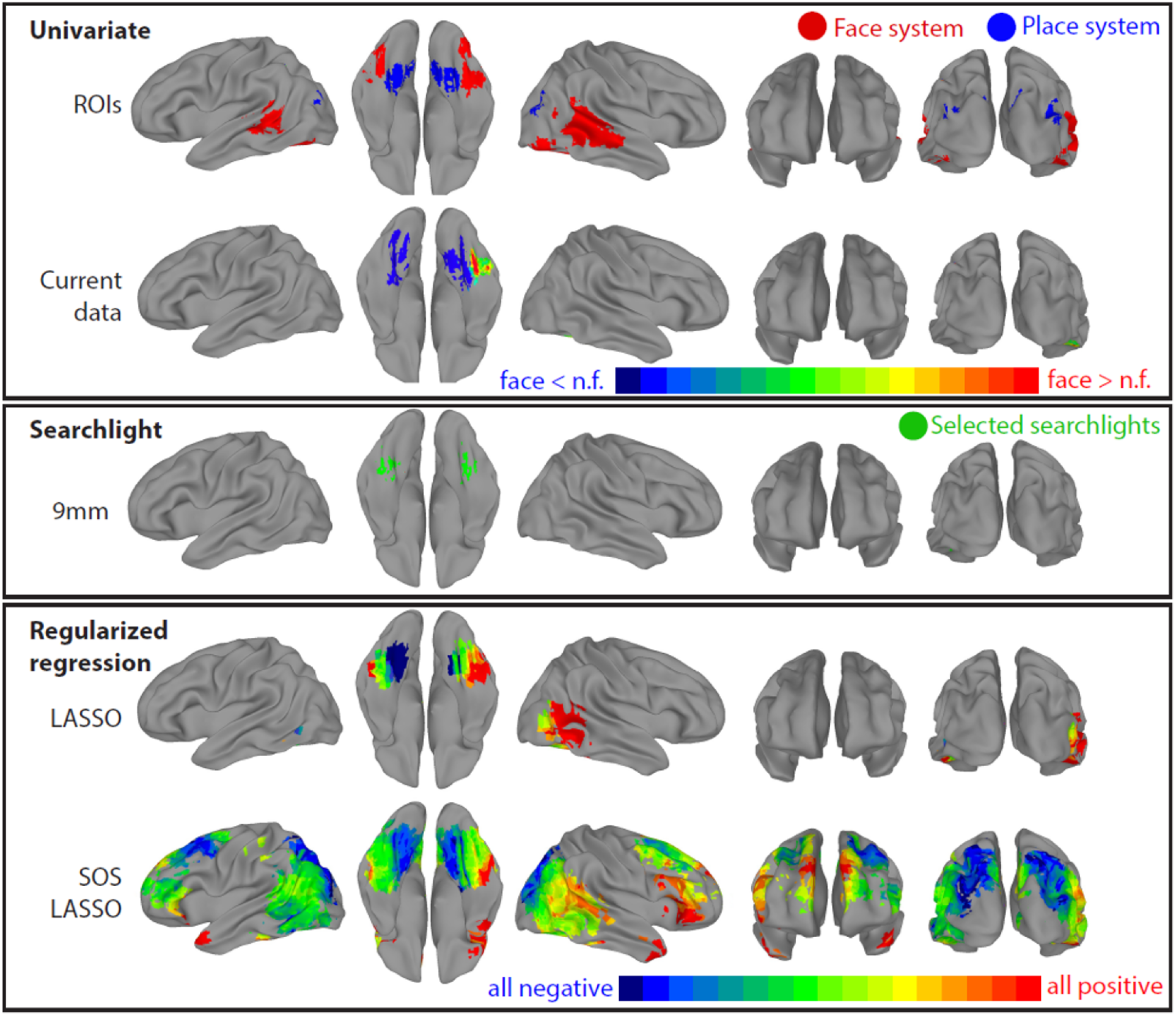
Regions discriminating faces from non-faces according to four approaches. *A. Univariate contrast results*. Top: the canonical “face” and “place” systems. Bottom: univariate contrast results for the current data. The bilateral non-face effect (blue) is statistically reliable in a cluster-corrected whole-brain analysis while the r¡ght-hemisphere face effect (warm colors) is reliable within the right fusiform face area. *B. Results from searchlight MVPC*, shown in green since the direction of the underlying neural code is undetermined in this approach. Colored regions are statistically reliable in a cluster-corrected whole-brain analysis *C. Results from whole-brain regularized regression*. Colored regions are those selected more often across subjects than expected under the null hypothesis according to permutation testing. Hue indicates the proportion of classifier coefficients that are positive across participants. Top: LASSO reveals a pattern largely consistent with the existing literature. Bottom: SOS-LASSO encompasses these regions along with a much broader distributed signal not revealed by the other methods.

Searchlight MVPC with a 9mm radius identified a localized signal in regions near the FFA bilaterally. LASSO selected a handful of voxels in each subject, concentrated near FFA bilaterally and in right lateral occipito-temporal cortex. The classifier coefficients indicated that reduced activity predicted faces in medial ventro-temporal cortex while elevated activity predicted faces for lateral ventro-temporal and occipito-temporal regions. Together the three approaches suggest somewhat different conclusions about face representation in cortex, each having precedent in prior work: the UC result suggests that faces selectively activate the right FFA;^28^ searchlight identifies localized but bilateral signal around the FFA;^29^ and LASSO reveals a nonface-to-face gradient bilaterally in these regions plus a right-lateralized occipital “face” region.^30^

Results from SOS LASSO differed strikingly. Colored regions in Fig. 3C (bottom) show voxels that received non-zero weights on significantly more subjects than expected from a null distribution estimated by permutation testing (p < 0.001 corrected; see Methods). Hue again indicates the proportion of selected coefficients that were positive across subjects. In addition to the canonical face and place systems the results implicate a host of regions spanning anterior temporal, frontal, and parietal cortex.

We assessed the quality of the representations discovered by each method as follows. For each subject and method, we trained a (ridge-regularized) logistic classifier to discriminate face from non-face stimuli using only the voxels that the method identified as important, then compared the classifier cross-validation error (mean of hit and correct rejection rates) across methods. All classifiers performed reliably better than chance (50%) but with substantial differences among them. Searchlight solutions performed worst (67%); the canonical univariate ROIs yielded significantly higher accuracy (76% correct, p < 0.001 vs searchlight); but performance for LASSO and SOS LASSO was even better (88% and 87% respectively, p < 0.001 vs univariate for both) and did not differ significantly from each other.

Why do neural representations appear so widely distributed under SOS LASSO? Perhaps the neural signal outside canonical areas merely serves to de-noise, interact with, or otherwise amplify the signal encoded within the canonical face and place systems (see SI-Denoising).^31^ We assessed this possibility with two additional analyses. First we fit SOS LASSO to two subsets of data: voxels lying within the canonical face and place systems (padded by an additional 7mm) or voxels lying outside these systems and also not selected by any other method. If the true signal lies mainly in the face and place systems, the within-system classifiers should yield higher accuracy on held-out items than outside-system classifiers. Instead the reverse result obtained: outside-system classifiers showed significantly higher hold-out accuracy (87%) than within-system classifiers (83%; t(11) = 2.4, p < 0.05), and as well as whole-brain classifiers (87%).

Second, we used SOS LASSO to find voxels that reliably discriminate place from non-place stimuli. If broadly-distributed regions are selected because they de-noise system-specific signal or for some other spurious reason, they should be selected regardless of the category decoded. Figure 4 shows this is not the case: the voxels that discriminate place from non-place stimuli are anatomically localized and consistent with the canonical view of place representation in the brain.^32^ Together the results suggest that widely distributed regions outside canonical systems contain information that helps to discriminate visually-presented faces from non-face stimuli.

**Figure 4.**
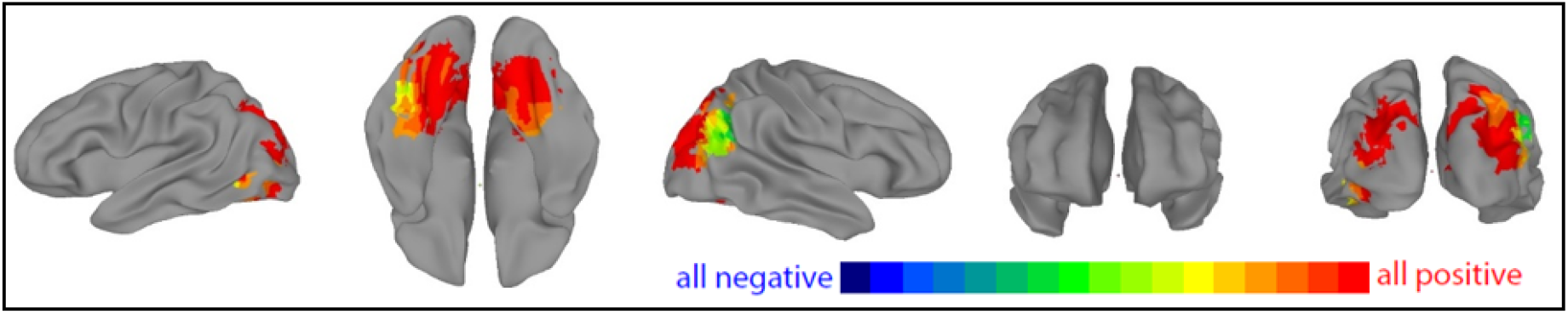
Regions discriminating place from non-place stimuli according to SOS LASSO. The results are consistent with standard views of scene representation in the brain, demonstrating that SOS LASSO does not always find widely-distributed solutions.

## General Discussion

Using computer simulations we have shown that four contemporary statistical methods for brain imaging have significant blind spots remedied by a new approach based on structured sparsity, the SOS LASSO. All methods yielded qualitatively different results when applied to fMRI data collected while participants judged visual images of faces, places, or objects, but solutions from prior approaches were generally consistent with the standard view that face representations are encoded within posterior temporal and occipital cortices. The SOS LASSO yielded a much more widely distributed solution encompassing anterior temporal, frontal, and parietal regions.

Can we be confident that SOS LASSO has revealed real signal? The result does not solely reflect selection of spurious voxels outside canonical systems, since (1) extra-system regions were sufficient to discriminate face from non-face stimuli with high accuracy and (2) the same approach yielded a more localized solution when seeking signal that discriminates place from non-place stimuli. Might it have arisen from idiosyncrasies of the particular stimuli or task used in the current study? We cannot rule the possibility out, but the current data were sufficient to replicate canonical results using prior methods—only with SOS LASSO was further signal revealed. Thus it remains possible that prior work has likewise missed extensive distributed signal.

Many aspects of the SOS LASSO solution accord well with both standard views of face and place perception and with the broader literature. Regions where classifier coefficients are consistently positive (warm colors in Figure 3) pick out much of the canonical face system and other cortical areas known to support social cognition, including the bilateral temporal poles,^33,34^ right orbito-frontal cortex,^35,36^ and superior medial-frontal cortex.^37^ Regions where coefficients are reliably negative (cool colors in Figure 3) pick out areas known to encode less socially-critical information, including scenes (parahippocampal place area)^32^ and object-directed action (dorsal visual stream, left dorsal premotor motor area)^38–40^. The mixed-direction regions (green in Figure 3) suggest that, in addition to these directionally-consistent codes, information about the stimulus category can be encoded with distributed patterns in which the direction of the code varies within and across individuals. The current results cohere with other recent work suggesting that neural representations may be more broadly distributed that heretofore suspected, for face perception specifically^41,42^ and for conceptual structure more generally.^43,44^

The pattern of results across methods has two important implications for hypotheses about neural representation. First, the out-of-system information detected by SOS LASSO was not discovered by even a large searchlight, indicating that it does not reside within local cortical regions considered independently. Thus *anatomically distal regions can jointly encode multivariate representational structure*. Second, SOS LASSO assigned consistently positive or consistently negative classifier weights in regions where univariate contrast yielded a null result. Whereas univariate contrast considers mean activations of each voxel independently, a classifier coefficient indicates the voxel’s contribution to the classification decision while taking the contributions of other voxels into account. Similar to partial correlation analyses, a voxel’s activity may not correlate reliably with stimulus category when considered alone, but may correlate with residual variation once the effects of other voxels have been factored in. In that case the voxel will receive a weight in the classifier but will not show a significant effect in univariate analysis. The contrasting results thus indicate that *voxels can contribute to representational structure in a directionally and locationally consistent manner even when they do not independently correlate with that structure*.

These observations suggest a new perspective on face representation in the brain. The standard view posits “core” and “extended” systems residing within posterior temporal and occipital cortices. ^45,46^ SOS LASSO suggests that, in addition to these regions, face perception generates widely-distributed activation patterns across much of the cortex, with anatomically distal regions jointly helping to differentiate the representations of visually presented faces from other stimuli. Within the full pattern, faces are signaled by elevated activity in social-cognitive brain networks, by suppressed activity in networks relevant to navigation and object-directed action, and by heterogeneous patterns in parietal and frontal cortices—but because elements of the pattern vary in the direction, independence, and localization of their code, previous methods each provide only a partial view of the full representation. Their agreement with the standard view arises because classic “core” and “extended” face systems comprise parts of the pattern where information is encoded efficiently and independently within circumscribed cortical regions, often in a directionally-consistent manner localized similarly across subjects.

Relation to other multivariate brain imaging approaches. We have focused on the common experimental design wherein the theorist seeks neural signal that differentiates one discrete experimental condition from another. Other designs have other goals and so employ other methods—for instance, representational similarity analysis (RSA) seeks voxel sets that jointly express a target similarity structure,^47^ while “generative” approaches seek to predict whole-brain images from externally-derived features of the stimulus or experimental condition.^48^ Like MVPC, such approaches often adopt techniques that can limit their ability to detect network-distributed signal, such as independent consideration of each voxel time-series or of different anatomical regions. Structured sparsity may likewise provide new insights for these kinds of problems.^49^ We also note that SOS-LASSO constitutes just one way of leveraging structured sparsity for pattern classification—other groups are pursuing similar ideas,^24,50^ and understanding the relations among these approaches is a central goal for future research.

### Broader implications for cognitive neuroscience

Our aim has been to consider how functional imaging results might change when one employs statistical methods capable of discovering distributed signal of the kind suggested by neural network models of cognition. We chose visual face perception as one extensively-studied domain where both univariate and multivariate functional imaging have highlighted fundamental questions about neuro-cognitive representation. The contrasting results from SOS-LASSO compared to other methods illustrates that such an approach can lead to quite different conclusions even in well-studied domains. Yet such a discrepancy is not guaranteed: the decoding of place information with this method yielded results highly consistent with canonical views. The current study thus raises the possibility that functional imaging has provided an incomplete picture of neuro-cognitive representation across domains—but assessing that possibility in any given domain will require further work of this kind.

## Online Methods

### Simulation study

#### Model implementation

Code and documentation for running the simulations and subsequent analyses appear at https://github.com/crcox/SOSLassoSimulations. The model shown in Figure 2A of the main paper was trained on 72 items sampled from two domains, A and B. Each item activated exactly 2 systematic and 2 arbitrary input units, and across items each unit was active in exactly 8 items. Half of the systematic units were activated only by items from domain A, while the remaining half were activated only by items from domain B. Thus any pair of items in the same domain had a small probability of overlapping in some of their systematic properties, while items from different domains never overlapped in their systematic properties. Arbitrary units were equally likely to be active for items from domain A versus B.

The model was fit using the Light Efficient Network Simulator (https://github.com/crcox/lens) using back-propagation to minimize cross-entropy error. The weights were adjusted with a learning rate of 0.1, using momentum (“Doug’s” momentum = 0.9) and subject to weight decay (decay constant = 0.001). The model was trained 10 times to asymptotic performance with very low error over 1000 epochs. Prior to each training run, the model was initialized with random weights sampled from a uniform distribution in the range [−1, 1]. These 10 models were used to generate data for 10 model “subjects,” based on the patterns of activity elicited by each input over the whole network. Each model was presented with the 72 input patterns in sequence, and the pattern of activation elicited over the 114 units in the network (including the 28 irrelevant units, which always had an activation of zero) was recorded. The dataset for each model subject thus consisted of a matrix with 72 rows corresponding to stimulus items and 114 columns corresponding to model voxels. Each matrix contained the “true” response pattern for each subject to each item. To simulate noise in the measurement of this activity, a random value sampled independently from a Gaussian distribution with a mean of zero and standard deviation of 1 was added to each cell of the matrix. We take the resulting values in each cell of a matrix to be a model analog of the estimated BOLD response to a single stimulus at a single voxel in a single subject in an fMRI study.

To apply different brain-imaging methods to the discovery of structure, it is necessary to further stipulate the anatomical locations of the different units the model. In all simulations, input units were situated all together, with domain-A units neighboring one another, domain-B units neighboring one another, arbitrary units neighboring one another, and the three units types concatenated end-to-end (AH units immediately adjacent to SH units; irrelevant units immediately adjacent to AH units). Output units were organized the same way, though outputs were assumed to be anatomically distal to inputs. The anatomical arrangement of input and output units was identical across model individuals.

For hidden units, we considered two different anatomical organizations. For *anatomically localized* models, units within a hidden layer (SH, AH, or irrelevant) were treated as anatomical neighbors with the unit types again concatenated end to end and localized in the same way across model individuals. Consequently, large searchlights or over-zealous smoothing could integrate activation from more than one hidden unit type. In the *anatomically dispersed* condition, hidden and irrelevant units were organized into three “regions” each containing a mix of SH, AH and I units, plus a fourth region containing only I units. The regions were each treated as anatomically distal to one another and to the input and output units. The three regions with mixed unit types all contained 7 units total, with 2 or 3 SH units, 2 or 3 AH units, and 2 or 3 I units. All model subjects had the same 7 units assigned to a given region, but the spatial ordering of these units within region was shuffled at random independently for each. The aim was to generate a scenario in which spatially distal units can jointly encode representational structure, with the coarse anatomical layout consistent but fine-grained layout variable across individuals. Note that model connectivity, the activation patterns evoked by different inputs, and the ways these patterns were distorted by measurement noise were identical for both model layouts—all that differed was the spatial locations of the units in each layer.

#### Statistical analysis

Each analysis ultimately involves computing, for each unit, the probability of the observed result under the null hypothesis that the unit does not contribute to differentiating A from B items. For fair comparison across methods, we therefore used the same statistical threshold for “counting” a unit as significant (p < 0.002 uncorrected). The qualitative pattern of results across methods does not vary with this criterion.

##### Univariate contrast

The activity at each unit was analyzed for all subjects using a mixed-effects model that treated subject as a random factor^1,2^, stimulus category (A or B) as the sole fixed effect, and unit activation as the dependend measure. The model was fit using the fitlme function in MATLAB, and the fixed-effect coefficient was tested for significance using the Satterthwaite approximation to the degrees of freedom and a standard F-test, numerator degrees of freedom = 1, denominator degrees of freedom = 9. The results are directly analogous to a repeated-measures ANOVA. Results were thresholded at uncorrected p < 0.002. The analysis was conducted for both the anatomically localized and the dispersed model. In both cases, the data were spatially smoothed, taking a weighted average over a three unit window, where the center unit was weighted about twice as much as the two flanking units.

##### Searchlight MVPC

The searchlight analysis was conducted using the SearchMight toolbox^3^ for MATLAB. Non-contiguous unit groups from Figure 2A in the main paper were treated as anatomically separated regions so that a searchlight never encompassed units in different regions. This was accomplished by inserting empty units between the layers, and providing a mask to SearchMight to omit those units during analysis. Within each searchlight, a Gaussian Naive Bayes (GNB) classifier was fit to distinguish between category A and B items. Although GNB classifiers are limited in some ways,^3^ the concerns do not apply to this simple and idealized case where noise is truly identically and independently distributed (iid) with uniform variance. Classifier performance at each searchlight was estimated through 6-fold cross validation. The mean cross-validation accuracy was stored at each searchlight center, and the mean accuracy over model subjects was tested for each unit to see if it differed significantly from chance. The resulting map of p-values was thresholded at p < 0.002 uncorrected. The analysis was performed on both the anatomically localized and the anatomically dispersed arrangements of units. The data were not smoothed prior to the searchlight analysis. The analysis was performed with small (3 unit), medium (7 unit), and large (14 unit) searchlights; results for smaller and larger searchlights are reported in SI-Model.

##### LASSO

Logistic LASSO (and ridge regression; see SI-Model) were conducted as special cases of the SOS-LASSO optimization code posted at https://github.com/crcox/WholeBrain_MVPA/blob/master/src/%4QSOSLasso/SOSLasso.m, which is part of the broader Wholebrain Imaging with Sparse Correlations (WISC) workflow maintained by CRC. Both methods have a free parameter *λ* that controls the importance of the regularization penalty relative to the prediction error, leading to greater sparsity in LASSO and more severe weight shrinkage in ridge regression. The analysis thus proceeded in two steps: one to estimate a useful *λ* for each simulated subject, and a second to fit a model at the estimated *λ* and evaluate it on a hold-out set. The data for each simulated subject was first divided into 6 equal parts, each containing the same number of category A and B items. One part was set aside and the remaining 5 were passed to a function that conducted a 5-fold cross validation accuracy search across many values of *λ*. The parameter search was implemented using hyperband, a state of the art search procedure that optimizes use of parallel computing infrastructure when fitting complex optimizations^4^. The function returns the *λ* producing the highest cross-validation accuracy, which is subsequently used to fit a model to all 5 parts of the data. The resulting model was then assessed on the original hold-out set (the 6th part). This procedure was carried out separately for all 10 model subjects, in both localized and anatomically dispersed model variants.

LASSO provides a straightforward selection criterion: any unit receiving a non-zero weight has been “selected” as important for predicting the stimulus class. We conducted a permutation test to assess, for each unit, the probability of selection under the null hypothesis of no relationships between unit activation and category label. We first estimated the best sparsity level *λ* from the unpermuted model data using the procedure just described. For each permutation, we randomly shuffled the category labels, then fit a LASSO model at the pre-specified *λ* and recorded the resulting coefficient on each unit. 1000 permutations were conducted. We then estimated the probability of selection for each unit as the number of times the unit received a non-zero weight divided by 1000. These probabilities were estimated separately for each model subject, then averaged across model subjects at each unit, yielding a base probability of selection for each unit. In the non-permuted data, we counted how often a given unit received a non-zero weight across the 10 simulated subjects, then used the binomial distribution to compute the probability of this outcome given the base probability of selection from the permutation testing. For instance, if a unit was selected with probability 0.2 in the permutation testing, and was selected in 5 out of 10 simulated subjects in the real (non-permuted) data, we computed the likelihood of this outcome under the null hypothesis as the binomial probability of achieving 5 or more successes in 10 attempts given a base probability of 0.2 for success (p ~= 0.033). This probability was computed for every unit and the results were thresholded at p < 0.002 without correction for multiple comparisons.

##### SOS LASSO

The SOS LASSO analysis was implemented using custom code built on top of MALSAR^5^ for MATLAB. The data were divided into overlapping groups based on “anatomical” proximity. Each set included 7 units (except for the last group in the input and output layer, which have 8 rather than having one unit on its own), and the groups did not overlap. The set size was selected to correspond with the number of systematic hidden units, to emphasize the difference between the localized and dispersed cases (i.e., group size was optimal for the localized condition, and ensured that systematic hidden units were maximally dispersed over groups in the dispersed condition). SOS LASSO has two free parameters, one controlling sparsity at the set level and one controlling overall sparsity. As in the previous analysis, these parameter values were selected through an internal 5-fold cross-validation process, then a final model was trained with the best parameters and tested on a sixth hold-out set.

To statistically determine which model units were reliably selected we again conducted a permutation test. The true category labels were permuted 1000 times and for each permutation we fit a SOS LASSO model to all 10 model subjects simultaneously, using hyper-parameters chosen from the true data. We then computed, for each unit, the proportion of times it was selected across all permutations and all model subjects, taking this as the probability of selection under the null hypothesis. The true data were thresholded using the binomial distribution to compute, for each unit, the probability of the observed number of selections out of 10 model subjects given the null-hypothesis base probability for that unit. We again thresholded this probability map at p < 0.002 uncorrected.

### Imaging study

#### Data collection methods

The fMRI dataset was collected and contributed by Lewis-Peacock and Postle.^6^ Subjects viewed 90 stimuli from three categories: 30 famous people, 30 famous locations, and 30 common objects. They then indicated how much they liked the celebrity, how much they would like to visit the location, or how often they encountered the object in everyday life using a stimulus-response box and a four-point Likert scale. Each stimulus was presented one time only, for a total of 90 randomly ordered stimulus presentations. Each trial consisted of a cue period (2 s), a stimulus period (5 s), and a judgment period (3 s). Each trial was followed by an arithmetic task (16 s) to reduce interference between trials.

Whole-brain images were acquired with a 3T scanner (Signa VH/I; GE Healthcare). T1-weighted images (30 axial slices, 0.9375 x 0.9375 x 4 mm) were acquired for all 10 subjects. Functional images were acquired using a gradient-echo, echo-planar sequence [repetition time (TR), 2000 ms; echo time, 50 ms] within a 64 x 64 matrix (30 axial slices coplanar with the T1 acquisition, 3.75 x 3.75 x 4 mm). Six scans were obtained for each subject, each scan lasting 6 min, 50 s. Each scan was preceded by 20 s of dummy pulses to achieve a steady state of tissue magnetization. Preprocessing of the functional data was done with the Analysis of Functional NeuroImages (AFNI) software package^7^ using the following preprocessing steps, in order: correction for slice time acquisition and rigid-body realignment to the first volume from the experimental task; removal of signal spikes; removal of the mean from each voxel and linear and quadratic trends from within each run; and correction for magnetic field inhomogeneities (using in-house software). Note that spatial smoothing was not imposed, and the data were not spatially transformed into a common atlas space before hypothesis testing. Rather, the data from each subject were analyzed in that subject’s un-smoothed, native space, without fitting a GLM to the hemodynamic response. Instead, the BOLD response at the 5th TR after stimulus onset was selected for analysis, which was selected to be near the anticipated peak of the hemodynamic response.

#### Statistical analysis

In all analyses the functional data for each subject were first masked to exclude voxels outside of cortex (white matter, CSF, cerebellum, thalamus and sub-cortical structures, bone, etc).

#### Univariate analysis

The functional data for each subject were projected to Talairach space using the T1 data with a combination of manual landmark identification (anterior and posterior commissures) and automated affine transformation obtained using 3dvolreg in AFNI. The spatially normalized response to each stimulus, sampled at the 5th TR after stimulus onset, was smoothed with 4 mm FWHM Gaussian kernel. At each voxel we computed the mean response for face stimuli and for non-face stimuli and subtracted these (face – nonface). In a whole-brain analysis, we computed a t-test on this difference against the null hypothesis of 0, and thresholded the resulting map using a cluster-corrected whole-brain threshold of p < 0.05. The resulting whole-brain map revealed bilteraral parahippocampal regions that were less active for faces than for non-face stimuli, but no regions that were more active for faces. We then conducted a region-of-interest analysis directly assessing whether the right fusiform face area (rFFA)—the earliest and most consistently reported face-selective area in cortex—was reliably more active for face than non-face stimuli. Using the rFFA mask published by Kanwisher’s group, we computed the mean difference in response for faces and for non-faces across all rFFA voxels in each subject, took the difference, and computed a t-test against the null hypothesis of 0. The rFFA was reliably more active for faces with p < 0.03, one-tailed. The same ROI analysis was then conducted for each other face-specific region in the canonical maps, but no other region showed a reliable difference in activation for faces versus non-faces.

#### Searchlight MVPC

We used SearchMight with a GNB classifier to generate native-space information maps for each subject using both a 9 mm and a 15 mm searchlight around each voxel. The maps were first normalized to Talairach space using the same procedure described for the univariate analysis, and then smoothed with the same 4 mm Gaussian kernel. The performance metric for each searchlight (i.e., the value stored at each point in the information map) was the difference between the hit rate and the false positive rate, which has an expected value of 0 under the null hypothesis. A one-sample, one-tailed t-test against 0 was conducted on this metric. P-values were thresholded at p < 0.05 using the same whole-brain cluster-correction approach employed with univariate analysis.

#### LASSO

The analyses were completed in two rounds of modeling, to support two different statistical analyses: one to test the generalization accuracy of the fitted models (the performance round), and the other to assess which voxels reliably contribute to that performance (the importance mapping round).

##### The performance round

Model performance was evaluated independently for each subject using cross-validated generalization error. The workflow for evaluating performance (Figure SI-1) proceeds as follows. The available examples (i.e., whole-brain maps showing the BOLD response generated by each item in the experiment) are split into 10 blocks; these block assignments are the same for all analyses reported in this document. Before model fitting, one block is set aside as the final holdout set, and model fitting then loops over the remaining nine blocks, holding out each one in turn as a test block and using the remaining 8 to “tune” the free hyperparameter. Many models are fit using the tuning data and different hyperparameter values and each such model is evaluated against the same test block in that loop. This yields a set of error values, denoted as *E*_*λ*1_…*E_λn_* in the diagram. The same hyperparameter values are used in each iteration of the loop, and the model prediction error associated with a given hyperparameter value is estimated as the mean error across all 9 loops. The hyper-parameter value that produces the lowest mean error is then used to fit a model to data from all 9 tuning blocks and this model is then evaluated on the final holdout set. The whole procedure is repeated 10 times, with each block in turn treated as the final holdout set. This yields 10 “final error” values, denoted as *E*_*f*1_…*E_fn_* in the diagram, which are averaged to produce an estimate of the classifier performance. This nested cross validation procedure ensures that the final holdout set is completely isolated from the hyper-parameter selection and model fitting stages, and also helps avoid idiosyncrasies of particular combinations of training and test sets.

**Figure SI-1:**
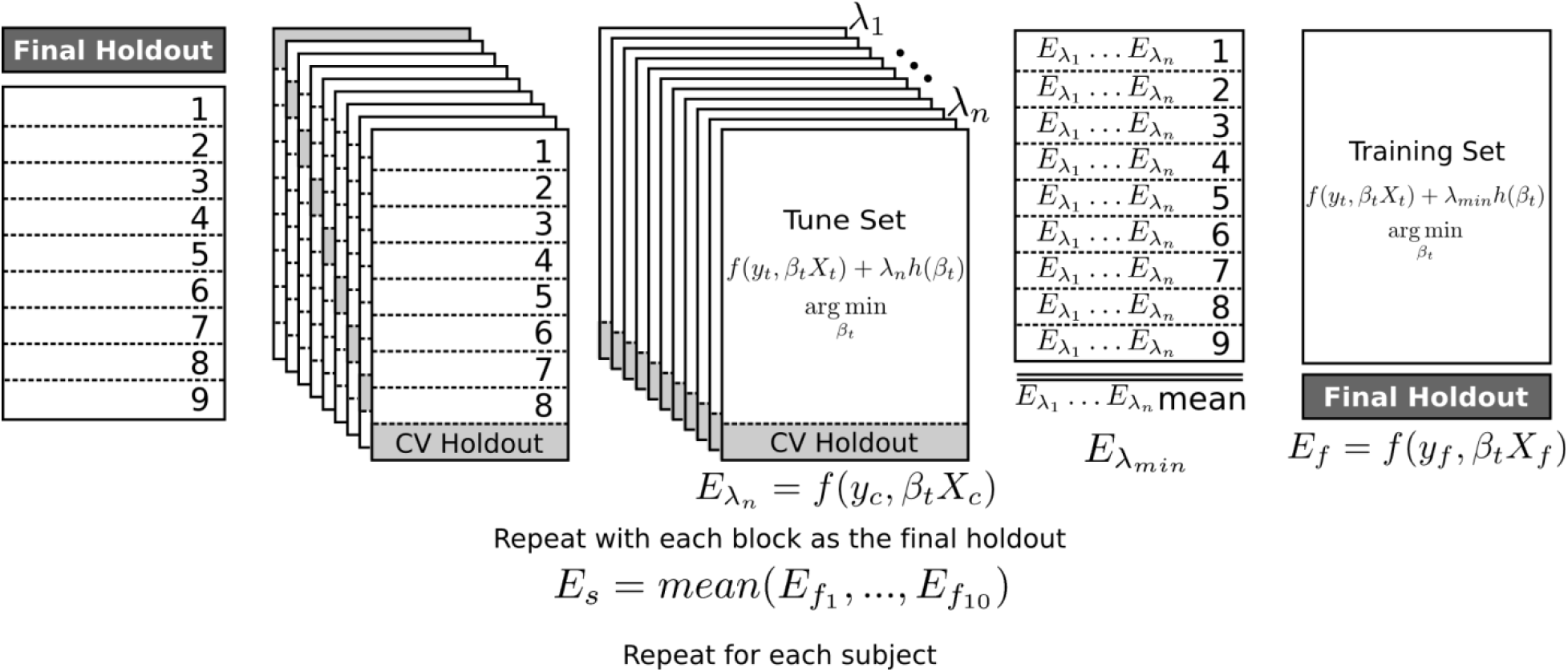
Nested cross-validation procedure for assessing model performance.

*The importance mapping round* fits a single model for each subject using all data, with the aim of then analyzing the model coefficients to determine which voxels are contributing in what way to classifier performance. The workflow for importance mapping (Figure SI-2) is similar to the prior round except that the outer loop over final holdout sets is eliminated so that a single model is fit to each subject using the entire dataset. As for the performance round, hyperparameters are tuned with 10-fold cross validation on model hold-out error. The best hyperparameters are then used to fit a final model to all data from all subjects. The coefficients from this model are then analyzed.

The motivation for the two steps is as follows. To get an accurate estimate of classifier performance, it is important that the final test items are excluded form all model-fitting steps, including hyperparameter selection, as done in the performance round. Several folds of validation must be conducted to ensure that the performance estimate is not unduly influenced by the particular items selected for the hold-out set—but this means that a different model is fit for each hold-out fold, and that the best set of hyperparameters may differ across folds. It is then unclear which model or hyper-parameter set should be used to interpret coefficients. Thus the performance round is conducted only to assess the expected hold-out error for the classifier. The importance mapping round does not provide useful information about expected hold-out error but yields a single model with one weight per voxel per subject.

To interpret these weights, the coordinates associated with each non-zero value were warped into Talairach space and the weight values at each point were linearly interpolated onto a common 3x3x3 mm grid. These interpolated weights were then smoothed with a 4 mm FWHM Gaussian kernel to compensate for imperfect warp alignment in each subject. To determine which voxels were selected across participants more often than expected under the null hypothesis, we fit 1000 permuted models for each subject, using the same functional data with the category labels shuffled randomly, and using the same hyperparameters selected from the properly-labeled data. The permutation solutions were projected into Talairach space and smoothed in the same way as the real data. For each voxel in each subject we computed the probability of selection (non-zero value) from these permutations. This allowed us to assess, for each voxel in each subject, its probability of selection using the data-tuned hyperparameters but with no systematic relationship between the fMRI data and the target labels. This information was used to construct a group-level binomial test at each voxel, where the probability of a voxel being selected N times out of 10 was assessed relative to the base rate of selection over the 1,000 permuted models. Voxels selected 3 or more times were associated with an uncorrected p < 0.002, and this is the threshold applied to the LASSO solutions.

**Figure SI-2:**
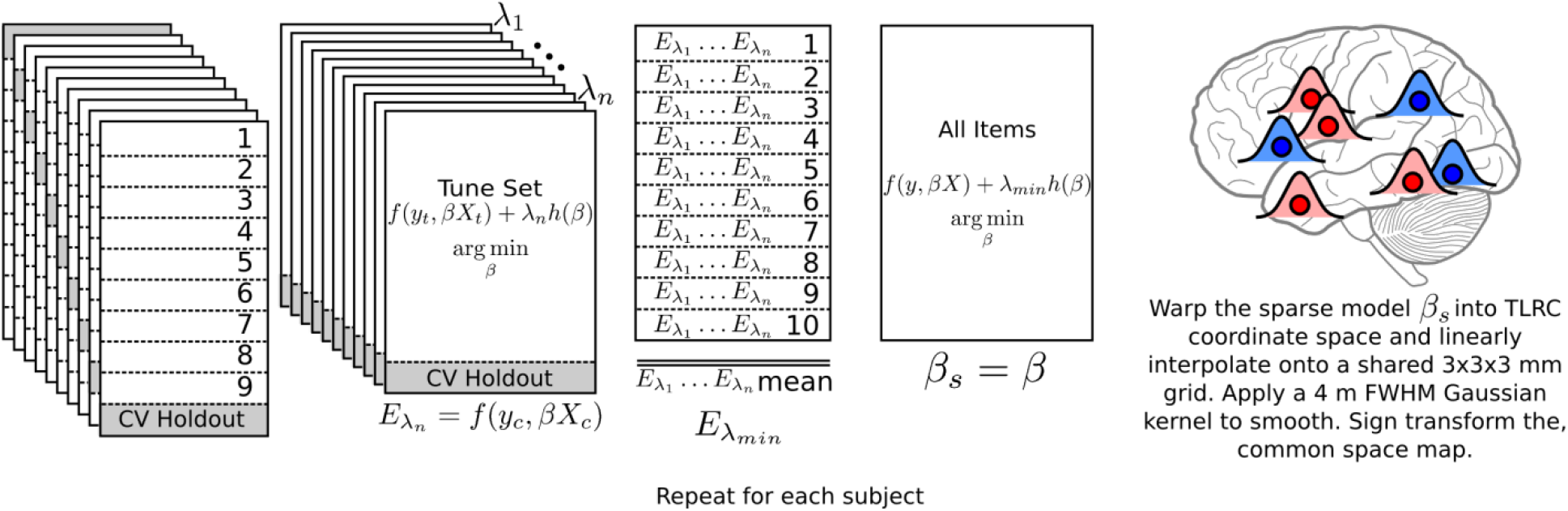
Workflow for the importance-mapping step.

#### SOS LASSO

As for LASSO, only cortical voxels were modeled, and analyses were completed in two phases. The performance phase proceeded with a nested cross-validation procedure exactly as described for LASSO. For the importance-mapping phase, appropriate hyperparameters were determined using 10-fold cross validation as with LASSO, and SOS LASSO solutions were then fit for 1000 permutations using these hyperparameters, shuffling the category labels in the same way for all participants on each permutation. Because the solutions are not independent across participants in SOS LASSO, we could not use the binomial approach developed for LASSO. Instead we counted, for each permutation, how often each voxel was selected across the 10 subjects. We then took the maximum of this value across all voxels and permutations—that is, the maximum across 1000 permutations * approximately 10,000 voxels. The largest amount of overlap observed across subjects in the permuted data was 7—we therefore thresholded the data map to show voxels selected in 8 or more of 10 participants. Because this level of overlap was never observed in the permuted analyses, we can be confident that the thresholded maps contain very few false alarms. To estimate the p-value, consider the number observations that went into determining the maximum amount of overlap in the permutation distribution, which is the number of voxels in the group map x the number of permutations. For 10,000 voxels, this would yield an uncorrected p < 0.0000001, which is lower than Bonferroni corrected p < 0.05 for 10,000 tests.

Because the SOS LASSO solutions implicated areas of the brain not often seen in analysis of face functional specificity, we performed follow-up analyses in which all voxels within the face and place systems published in ^8^ and padded by 7mm, or identified in any of the univariate, searchlight, or LASSO analyses, were excluded from the outset, leaving only the unexpected areas. We also ran the analysis only on voxels included within the padded face and place systems. Both analyses were conducted exactly as reported for the whole-brain analysis; results appear in the main paper text.

## Supplementary Information

### SI-NN: A brief overview of neural network models

Neural network models are composed of simple processing units that communicate via weighted synapse-like connections ^9,10^. Each unit adopts an activation state, typically varying between 0 and 1, that can be viewed as analogous in some respects to the mean firing rate of a population of spiking neurons proportional to their maximal rate.^11^ Units transmit information about their current activation through weighted connections, which can be viewed as capturing the net effect of activity in one population of neurons on another. Weights are typically real-valued, with negative numbers indicating a net inhibitory effect and positive numbers indicating a net excitatory effect. Each unit computes a simple process: it adjusts its current activation state according to the input it receives from other units in the networks. Input is usually computed as the inner product of the activation across all sending units and the values of the weights projecting from the sending units to the receiving units. The unit then converts the net input into a new activation state according to a specified transfer function (typically sigmoid, radial basis, or rectified linear). Units are conceived as computing inputs and updating activation states in parallel in continuous real time, though on serial computers this parallel process is simulated by updating units in discrete steps in randomly permuted order.

Within a network, units are organized into *layers* that govern the overall connectivity of the network: units within a layer tend to receive connections from, and direct connections toward, a similar set of units elsewhere in the network. Typically a subset of the units are specified to receive inputs directly from sensory systems (or other input systems outside the model), and to direct outputs toward motor systems (or other output systems outside the model). These unit subsets encode the input provided to the model and the outputs that simulate the model response. They are often referred to as *visible* units, because the theorist directly stipulates how different stimulus events and behaviors are represented with patterns of activation over the input and output units. Many models also include sets of units whose inputs and outputs are directed only to other units contained within the model—they do not receive external inputs from or direct outputs toward the model environment. For these *hidden* units, the theorist does not stipulate how different stimulus events or behaviors are to be coded with patterns of activation. Instead, the patterns of activation that arise across these units are determined solely by the values of the interconnecting weights. Deep neural network models are models with multiple layers of hidden units mediating between inputs and outputs.

The weights connecting units are shaped by learning and experience. Many different learning algorithms have been explored in this framework^12^, but all share the general idea that the weights gradually change over time in order to optimize some objective—for instance, minimizing the discrepancy between the outputs the model generates and the correct “target” outputs—as the network processes information from different stimulus events. Because the weights adapt to experience—that is, the structure of input and output patterns—and because the patterns of activation over hidden units depend upon the weight values, neural network models effectively acquire learned internal representations: the patterns of activation generated over hidden units by a given stimulus after the network has been trained. Neural network models can acquire internal representations that may initially seem counter-intuitive but can be shown, through computer simulations, to support behaviors documented in the domain of interest. One reason deep networks have proven useful in both cognitive science and machine learning is that they automatically acquire internal representations that are not intuitive a priori and so are difficult for the theorist to program or foresee.

Because network models abstract the neural processes they aim to uncover, they necessarily gloss many aspects of the structure and behavior of real nervous systems. Whereas individual neurons exhibit all-or-nothing spiking behavior, network units assume continuous activation states. Low-level dynamics such as lateral inhibition, temporal coherence, and local extra-cellular conditions are glossed in most connectionist models, while morphological differences among neuron types, cytoarchitecture, and other facts about brains are completely abstracted away. Network units are instead viewed as capturing, in a comparatively modest number of processing elements, the same informational states existing across vast numbers of heterogeneous spiking neurons in real nervous systems^10,13^. The central assumption is that the representational content and cognitive functions expressed in the coordinated spiking behaviors of hundreds or thousands of neurons behaving at a millisecond timescale can be usefully approximated by a much smaller vector of continuous-valued activations changing at a considerably slower rate.

The neural plausibility of network models has sometimes been challenged on these grounds, but it is important to note that essentially the same assumption is adopted by fMRI and other brain imaging methods. Such methods summarize the dynamical activity over hundreds of thousands of neurons behaving at a millisecond timescale with a much smaller vector of real-valued numbers, each expressing the overall metabolic demands exerted by populations of neurons averaged over some much coarser spatial and/or temporal extent^14^. The effort to relate cognitive events to neural activity so measured—that is, functional brain imaging—entails the assumption that important informational states over vast sets of rapidly time-varying neurons are preserved in smaller, coarser vectors capturing information about mean firing rates in local neural populations. In this sense, there is a natural affinity in the assumptions that neural networks and functional brain imaging methods adopt about how information is encoded in neural states.

In the current work, we therefore view the activation of a single unit in a network model as capturing the mean neural activity in a population of hundreds of neighboring neurons within a small volume of cortex as estimated at a single voxel from BOLD activity in fMRI. In the reported simulations we treat the activation of a single model unit by a given input pattern as analogous to the evoked response to a single stimulus item estimated at a single voxel in an fMRI study in a sparse event related design.^15,16^

### SI-Approaches. Conceptual overview of the different statistical approaches

Each statistical approach we consider deals with the complexities of brain imaging data by adopting important assumptions about how cognitive information is encoded in neural activity. The assumptions strongly constrain the kinds of signal that the method can discover, but these constraints are not always made explicit. This section provides a brief overview of the logic underlying each method and explicitly notes its core assumptions, summarized in Table 1.

**Table 1:** Assumptions implicitly adopted by different statistical methods for image analysis.

| Assumption | Description | Example | Univariate | Searchlight | LASSO | Ridge | SOSLASSO |
| --- | --- | --- | --- | --- | --- | --- | --- |
| Consistency of representation within ensemble | All units in a representational ensemble respond to objects of representation in the same way | Within an individual, all units representing faces will show greater mean activity for faces than for a control condition | Yes | No | No | No | No |
| Consistency of representation across individuals | A given unit within a particular representation responds to various stimuli in the same way across individual. | A unit that represents faces with greater mean activity in one individual will also represent faces with greater mean activity in other individuals. | Yes | No | No | No | No |
| Localization of representation within individuals | In any individual, anatomically neighboring units likely contribute to the same representation | Within an individual, a unit that shows systematically different responses for faces vs other objects will have neighbors that also show systematically different responses for faces vs other objects. | Yes | Yes | No | No | Partly: The optimization prefers solutions where useful features are anatomically proximal within individuals |
| Localization of representation across individuals | Across individuals, the units that contribute to a given representation will be found in similar anatomical locations | Though the units that encode faces may be distributed anatomically within an individual, the same distributed regions in other individuals are also likely to contain units that encode face representations. | Yes | Yes | No | No | Partly: The optimization prefers solutions where useful features are in similar locations across individuals |
| Voxel independence | The information coded by a unit activation does not differ depending on the activations of other units. | Units that represent faces do not "switch" to representing something else when other units are active. | Yes | No | No | No | No |
| Sparsity | Of all units measured, only a small proportion will be involved in the representation of interest | Only a small proportion of all units measured will be involved in representing faces. | No | No | Yes | No | Yes |
| Redundancy | The responses of units within a representation are highly inter-correlated | Within an individual, the pattern of responses over items will be highly correlated for all units involved in face representation | No | No | No | Yes | No |

*Univariate contrast analysis* has long been the standard method for interrogating fMRI data. It aims to identify regions of cortex that, across subjects, exhibit systematically different mean BOLD responses to two (or more) different kinds of cognitive events. Typically the BOLD signal is spatially smoothed: the raw response at each voxel is replaced with a weighted average of the responses from anatomically neighboring voxels. The smoothed time-series is then modeled independently at each voxel for each subject using a deconvolution procedure. This yields a beta coefficient for each experiment condition at each voxel indicating how well the measured BOLD signal matches the response expected if the activation of neurons within the voxel varies systematically with the experiment condition. The beta coefficients for each subject are projected into a common anatomical reference space, and univariate statistical tests are computed at each voxel independently to assess whether the coefficients differ reliably in the two experimental conditions across subjects. Voxels that show significantly different responses across subjects are viewed as important for coding the representation of interest.

A major challenge for the approach lies in establishing a meaningful criterion of significance in the context of tens or even hundreds of thousands of individual statistical tests. To avoid both false-positives and punishing corrections for multiple comparisons, it is common to seek ways of reducing the number of tests performed. Several different methods have been employed, but all rely on the idea that the representations of interest can be localized to particular cortical regions, and that the responses of voxels within a functional region will be largely similar. With these assumptions, the number of tests can be reduced by (1) conducting regions-of-interest analyses, where the responses of voxels within a ROI are averaged and the test is performed on the result mean response, (2) applying cluster-thresholding, where tests are only performed on clusters of *n* anatomically contiguous voxels all showing a similar response across subjects, or (3) applying a topographic control of the false-discovery rate. The univariate contrast method thus favors the discovery of clusters of anatomically neighboring voxels located in similar regions across individuals and showing similar response profiles across experimental conditions. From this brief description we can see that the method relies on five assumptions about the nature of the neuro-cognitive representations, summarized in the “univariate” column of Table 1.

#### Multivariate pattern classification

The remaining methods we consider are all variants of multi-voxel pattern classification (MVPC)^17^. Such approaches reverse the objective underlying univariate analyses: rather than using knowledge of the experimental design to explain variance in neural activity at individual voxels, MVPC uses the variance of neural activity across many voxels to make predictions about the experimental condition to which each trial, stimulus, or time point (henceforth, “example”) belongs^18^. To accomplish this, a classification algorithm is applied to a set of *training data* which include (a) the pattern of estimated activation evoked over a set of voxels for each of many examples and (b) a set of *labels* indicating the experimental condition or class associated with each pattern. For instance, in our model experiment, items in condition A might be labeled with a 0 while items from domain B are labeled with a 1. From the training data, the algorithm returns a pattern classifier—a statistical model that can be used to predict the label associated with any pattern of activation over voxels. Many different classification algorithms exist in the literature; as just one example, a logistic classifier will return a set of *weights*, one for each voxel, such that the estimated voxel activation, multiplied by its weight, summed over all voxels, and subject to a transformation function, yields a number that indicates the pattern label. In our example, a good logistic classifier should yield a number near 1 for all condition A items and near zero for all condition B items.

Even if there is no real signal at all in the data, it may be possible for a classifier to generate correct predictions for all items in the training set, especially when there are many predictors. Training set performance thus does not indicate whether the classifier is exploiting real signal in the data. Instead, the classifier is typically assessed on a *hold-out set*: an additional set of examples and labels collected in the same experiment but excluded from the training data. The classifier learned from the training data is applied to patterns in the hold-out set, and for each pattern it generates a “guess” about the associated condition label. The classifier output is compared to the true label to get a measure of accuracy. If a model performs above chance at classifying the hold-out set, this indicates that it is likely exploiting real information in the data. To ensure that the results do not depend upon the particular items chosen for the training and hold-out sets, it is common to test a model using *n-fold cross-validation*. On each “fold” a subset of items is chosen for the hold-out set, and different hold-out sets are selected for different folds, such that, across folds, all items appear in exactly one hold-out set. Each hold-out set provides a measure of model classification accuracy, and this is usually averaged across folds to provide a single number indicating how accurately the trained model can classify hold-out patterns. We will refer to this number as the *cross-validation accuracy* of the classifier.

MVPC algorithms, like univariate analyses, are challenged by the abundance of data provided by fMRI, and so must adopt additional assumptions about the nature of the underlying signal. In any fMRI study (as in our model) there will always be more predictors (voxels) than things predicted (stimulus items or events), producing an over-fitting problem. In such cases, there exists no unique solution to the classification problem defined by the training set. Closed-form analyses are undefined, and other model-estimation procedures will produce a classifier that perfectly fits the training data without any guarantee of finding real signal. These problems can only be addressed by constraining the analysis based on an underlying hypothesis about how signal is truly encoded in the data. As with the univariate method, these constraints systematically affect the results. Each of the remaining methods adopt different constraints to solve the over-fitting problem.

#### Searchlight MVPC

We begin with the well-known “searchlight” approach^19^, which was formulated specifically to address the challenge of finding distributed representations in brain imaging data. The method works as follows. Instead of training a classifier using all predictors at once, a separate classifier is trained for every individual voxel location in every individual subject. For each location, all voxels within a radius ***r*** of the center voxel are included as predictors in the classifier. This avoids the over-fitting problem by restricting the number of predictors included in any given classifier. The mean cross-validation accuracy for each classifier is stored in the searchlight center voxel, providing an *information map* for each subject. A univariate group-level analysis can be conducted on the information maps, similar in all respects to the analysis described in the previous section. Per the univariate assumptions, this means that each point in the accuracy map is considered independent of all others. However, within each searchlight, the effect of a given unit on the classification can differ depending upon the activations of other units in the searchlight, and these units can respond to various stimuli in quite different ways. Thus the searchlight method relaxes the assumptions that neural codes are consistent within and across individuals and that representational units are independent, but retains assumptions about localization of information within and across individuals (see Table 1).

#### Regularized logistic regression

Searchlights solve the over-fitting problem by restricting the field of view to a small number of contiguous voxels, then fitting many classifiers. An alternative is to train a single pattern classifier on all voxels at once. We used logistic regression as the classification model as this approach is easily interpretable, powerful, and draws upon intuitions formed through experience with linear regression.

A regression model is composed of a set of weights *β_x_*, one for each predictor variable *x* plus an additional intercept term, tuned to make accurate predictions about a response variable *y*. In logistic MVPC, the predictor variables are the voxels, and the response is a binary variable that codes class or condition label. For instance, in a contrast of conditions A and B, A events are labeled with *y* = 1 and B events are labeled with *y* = 0. To generate a prediction for a given item, the logistic regression model takes the weighted sum of the estimated response over voxels and passes it through a squashing function bounded at 0 and 1:

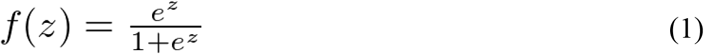

where:

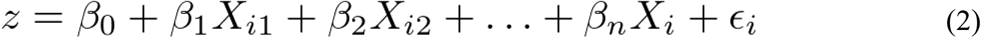

which is the model’s linear response to a particular pattern of activity. Thus, *f*(*z*) is a transformation of the weighted sum of predictor values expressing the probability that y=1 given the pattern of activity for the *i^th^* item. Fitting a logistic regression model involves finding coefficients that minimize the discrepancy between the true labels in *y* ∈ {0,1} and the probabilities assigned by the model. This is typically measured by the logistic loss:

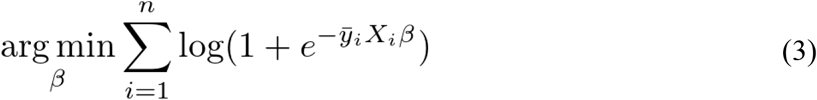

where *ȳ_l_* is −1 when *y_i_* = 0 and +1 when *y_i_* = 1. This loss is minimized when the sign of the model’s linear response *X_i_β* is positive for items labeled *y* = 1 and negative for items labeled *y* = 0.

The problem is that there are infinitely many solutions that will minimize this loss function when there are more predictors than items. One needs a way of deciding which among these is most likely to uncover the real signal. *Regularized* regression provides one way of doing this. Such approaches seek to jointly minimize the prediction error plus an additional cost, itself a function of the coefficients:

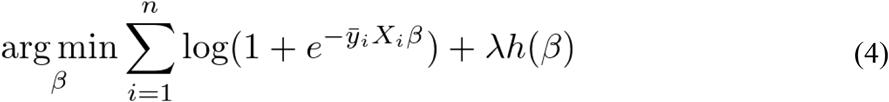

The additional penalty or *regularizer* represented by *h*(*β*) prioritizes some model solutions over others, and in this way embodies a hypothesis about the nature of the true underlying signal. The constant *λ* is a free parameter that controls the degree to which the two terms (prediction error versus minimization of the regularizer) should be weighted in the joint optimization.

In the main text we consider two varieties of regularized logistic regression employed in the fMRI literature: LASSO^20,21^ and *ridge regression*. ^22,23^ Though superficially similar, the two methods embody different implicit assumptions about the nature of the underlying signal and so yield quite different results. In LASSO, the regularizer is the sum of the absolute values of the model coefficients:

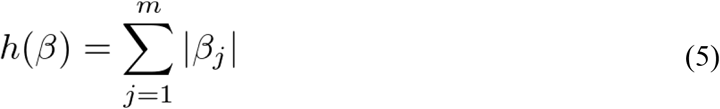

For ridge regression, the penalty is the sum of their squared values:

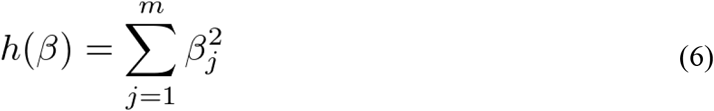

In both cases, the optimization is convex: for a given value of *λ*, there exists a unique set of coefficients that minimize the cost and that can be efficiently discovered by gradient descent. Yet the different penalties lead to quite different solutions. To understand why, it is useful to consider how they treat sets of predictors that covary together with one another. Imagine four voxels whose responses across items are perfectly correlated, and suppose their activations are useful in predicting the condition label. In this scenario, there are many different ways of placing weights over the four voxels that will all have the same effect on the classifier output. For instance, placing a weight of 1 on each voxel will have exactly the same effect as placing a weight of 4 on one voxel and a weight of 0 on the other three. Because the voxel activations are perfectly correlated, and the classifier operates on a weighted sum over voxel activations, these different weight configurations have the same effect on the model output and hence on the prediction error. The regularization penalty, however, should prefer some weight configurations over others.

If the data really are perfectly correlated, the LASSO penalty won’t be any help: the sum of the absolute value of the coefficients is the same for models that place a 1 on each unit versus models that put a 4 on one unit and zeroes on the rest. If we imagine, however, that all measurements are subject to some independent noise, the scenario is a bit different. In this case, one of the 4 units will, just by chance, covary slightly better with the category labels. In this case, the classifier can do a slightly better job of minimizing the error term by loading up all of the weight on this single voxel. Thus the joint optimization will lead to a solution where just one (or perhaps a few) of the redundant voxels are selected.

Ridge regression behaves very differently. Here the penalty scales exponentially as weights increase on a single voxel, but only linearly as weights are added across voxels. Thus the penalty is minimized by placing small weights on many voxels. In the preceding example, placing a weight of 4 on one unit and 0 on the remaining three leads to a total penalty of 16 over the four units. Placing a weight of 1 on each unit, in contrast, leads to a penalty of 4. Ridge regression thus prefers solutions where small weights are “spread out” over redundant predictors. In fact, as the weight approaches zero, the ridge penalty becomes vanishingly small, so with a finite number of training examples, ridge regression will always place at least a tiny weight on every predictor. In real data, of course, voxel states are never perfectly correlated nor perfectly informative about the condition label, so the behaviors of the two approaches are less easy to intuit. In general, however, it is useful to think of LASSO as minimizing prediction error with the *fewest* possible predictors (i.e., as many zero coefficients as possible), while ridge regression can be viewed as “spreading” small weights over all predictors exhibiting any systematic relationship with the category labels, without care for the number of predictors.

Table 1 summarizes the assumptions made by these approaches—all are relaxed relative to the univariate and searchlight methods. This does not mean that they are assumption-free. To the contrary, each approach entails additional assumptions. LASSO assumes the signal to be *sparse*, in that only a small set of uncorrelated voxels are involved in coding the information of interest. In this case, the best approach to finding true signal is to minimize prediction error using the smallest number of predictors possible. Ridge regression makes no sparsity assumption, but instead assumes that the signal is highly *redundant*, with many voxels expressing essentially the same information. In this case the best approach to finding true signal is to minimize prediction error using distributions of weights that are as close to zero as possible, so that all informative predictors are included in the solution.

### SI-SOS: The sparse-overlapping-sets (SOS) LASSO

The multivariate methods we have considered so far lie at two poles with regard to their assumptions. The searchlight method assumes consistent localization of signal within and across individuals. Regularized regression discards localization assumptions, allowing for the possibility that representations of interest are arbitrarily situated within and across individuals. The former assumptions seem too restrictive, the latter too loose. If human beings share the same gross neuroanatomical structure, then it seems reasonable to suppose *some degree* of consistent localization within and across individuals. Within individuals we might assume that, if a given voxel is important for representation, there is a good chance that some other units in the general vicinity are also important. Such an assumption could be adopted without further requiring that *all* neighboring units are important (univariate assumption 1), or that *only* neighboring units contain sufficient information for the representation (searchlight assumption 1). Likewise we might assume that, if a set of voxels contribute to a representation in individual A, then signal-carrying voxels are likely to be found in the same general anatomical vicinity in individual B. Such an assumption could be adopted without requiring that the location of the interesting information is very precisely aligned, or that the location contains useful information in all (or in the great majority of) individuals (assumption 3 for univariate and searchlight).

A central contribution of the main paper is to show how these intuitions can be formalized in a regularizer that promotes desired sparsity patterns amongst the classifier coefficients. The specific approach we describe is the Sparse Overlapping Sets (SOS) LASSO^24^. It adopts a central and intuitive assumption from *multi-task learning*^25^: if a set of learning tasks are related, then their solutions will also be related. In this case, each “task” is to find the neural activation pattern associated with a particular mental state in a single individual. In contrast to LASSO, we assume that the solutions across individuals are related: finding useful information in one individual gives us clues about where to look in another. The loose assumptions about localization within and across individuals can be built into a single optimization function that considers all subjects at once.

To achieve this, the voxels in the datasets from every individual are assigned to *sets*. Within an individual, voxels belonging to a given set are related to one another in some way--for instance, if the sets are defined by anatomical proximity, voxels within a set will all be near one another. Across individuals sets are aligned, so that voxels in *Set 1* for individual 1 are related to voxels in *Set 1* of individuals 2 through n. If sets are defined by anatomical proximity, then voxels in Set 1 will be localized similarly across subjects. The optimization then operates over all subject datasets simultaneously, minimizing an objective function that prefers solutions in which selected voxels belong to as few sets as possible. The end result is a solution that is unique for each dataset (i.e., each individual), but with the selected voxels tending to belong to the same sets within and across individuals.

Before considering the objective function itself, it is useful to consider how voxels are assigned to sets. Such assignments are made *a priori* and can in principle be guided by any hypothesis about how voxel sets are best grouped. For instance, sets might be determined by an anatomical segmentation, a previous analysis, a literature review, or any other prior knowledge. Importantly, however, set assignment can also be done in a relatively theory-neutral way: a volume can be split into a collection of *overlapping* sets of fixed size and degree of overlap, with each voxel belonging to multiple overlapping sets. On this approach, the voxels for each individual are first projected into a common anatomical reference space (without blurring). The space is divided into a 3-dimensional grid and, for each grid-point, all voxels within radius ***r*** are assigned to a given set. Each set thus contains a group of voxels in roughly the same anatomical location across subjects; each voxel belongs to multiple sets; and the sets overlap in the voxels they contain. The sets are analogous to searchlights, but rather than training separate classifiers for each set in each subject, a regularized logistic classifier is trained on all the data together. Because the optimization prefers solutions in which voxels belong to a small number of sets, such an approach will automatically select the particular grouping of voxels within and across individuals that best minimizes the objective. But, because the solutions are unique to each individual, there is no need for all voxels within a set to be selected or to receive similar weightings within or across individuals.

With that conceptual grounding, we can consider the SOS Lasso regularization penalty:

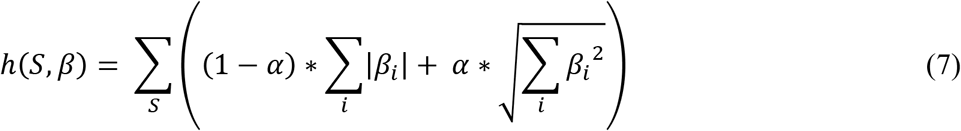

The function has two terms, Σ|*β_i_*| and 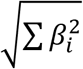, which are embedded in a loop that sums over sets. The first term is simply the Lasso penalty from Equation 5, computed separately for each set and summed over all sets. The second term, which we will refer to as the *grouping penalty*, takes the root of the squared coefficients summed over units within each set. *α* is a mixing parameter that determines the weight given to each of these two terms. When *α* = 0, the set penalty is not applied, and SOS LASSO effectively reduces to LASSO. When *α* is non-zero, however, the penalty behaves differently, preferring solutions where the non-zero coefficients (i.e., the selected voxels) all appear in the same overlapping sets—that is, when they tend to be in similar anatomical locations.

To understand how this preference is realized, we can compute the penalty associated with two competing solutions in a very simple case. Consider a 1-dimensional brain with 3 voxels. Voxels 1 and 2 are assigned to *Set 1*, and voxels 2 and 3 are assigned to *Set 2*. In the first solution, let voxels 1 and 3 be assigned a coefficient of +1, and voxel 2 a value of 0. The set penalty 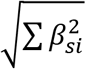 evaluates simply to 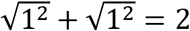, because the voxels are in different sets. In a second solution, let voxels 1 and 2 be assigned a coefficient of +1 and voxel 3 a value of 0. This time, the penalty evaluates to 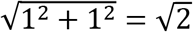, because the voxels are in the same set. Because the set penalty takes the root of the sum of squares across voxels in the set, the assignment of a +1 value to 2 of the 3 voxels produces a larger cost when the coefficients are placed on voxels from different sets.

There is a wrinkle here, of course: voxel 2 also exists in *Set 2*. If, however, we consider the coefficient assigned to voxel 2 as part of *Set 2*, it “costs” 1 penalty unit. If we consider it as part of *Set 1* it only costs 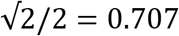 penalty units. In principle the total weight on the voxel could be apportioned across its corresponding sets in any manner; however allocation of the voxel’s weight to an otherwise empty group incurs a greater overall penalty, so the optimization ends up placing all of the voxel’s weights within sets that also contain other non-zero voxels. In this way the optimization “knows” to account for voxel 2 as belonging to *Set 1* rather than *Set 2*, and does not redundantly penalize the same weight. The further consequence is that the optimization automatically finds sets that contain many signal-carrying voxels if these are present in the data.

Finally, it is important to note that both the grouping penalty and the LASSO penalty increase when the magnitude of any weight is increased. This means that, just as with LASSO, solutions that include fewer voxels overall will tend to be preferred. Solutions will be sparse, even within groups. Sets are not selected themselves, but rather voxels are selected from within sets and it is less costly to select multiple voxels from the same set than from different sets.

In sum, the optimization prefers solutions that (a) fit the training data, (b) include units that are anatomically situated in similar sets, and (c) include a relatively small proportion of the units overall. As with LASSO and ridge regression, there is a tuning parameter that controls the weight given to the regularization penalty versus the error term; but there is also a second tuning parameter (*α* in the above equation) that controls the relative weighting given to the set penalty versus the LASSO penalty. A mathematical analysis of SOS LASSO and some applications of its use were recently described in ^26^ for linear regression and in ^24^ for logistic regression. The representational assumptions of SOS LASSO are summarized in Table 1.

### SI-Model

We here report some additional analyses of the simulation results.

#### Effects of searchlight size

Figure 2 of the main paper shows simulation results for a searchlight spanning 7 units, the best-performing searchlight size. When hidden units were localized consistently across model subjects, this approach excelled at discovering them and also did well identifying the signal-carrying I/O units. The approach failed to detect dispersed hidden signal, however. To assess how well searchlight analysis fares when the searchlight is not tailored to the size of the signal-carrying hidden units, we conducted the same analyses with smaller (radius 3) and larger (radius 14) searchlights. Figure SI-3 shows the results. Smaller searchlights reliably detected the systematic hidden units when they were localized, but missed the informative I/O units. The I/O units individually encode the category distinction very weakly, so small searchlights do not span enough of them to reliably decode the signal. The larger searchlights did a good job discriminating signal-carrying from arbitrary I/O units, but either completely missed (in the dispersed case) or seriously mischaracterized the location (in the localized case) of signal amongst the hidden units. A large searchlight centered on an uninformative unit can still perform well by virtue of including signal-carrying units in its periphery, mislocalizing the source of the good performance by storing it at the (uninformative) searchlight center. In this case, the searchlight applied to the localized layout ends up identifying all the *arbitrary* hidden units while missing the actual signal-carrying units. As a result of this phenomenon, the measure of accuracy—hit rate minus false alarm rate—was worse in the hidden units for the larger searchlight size even when hidden units were localized. When dispersed, no searchlight was able to discover useful hidden signal.

**Figure SI-3:**
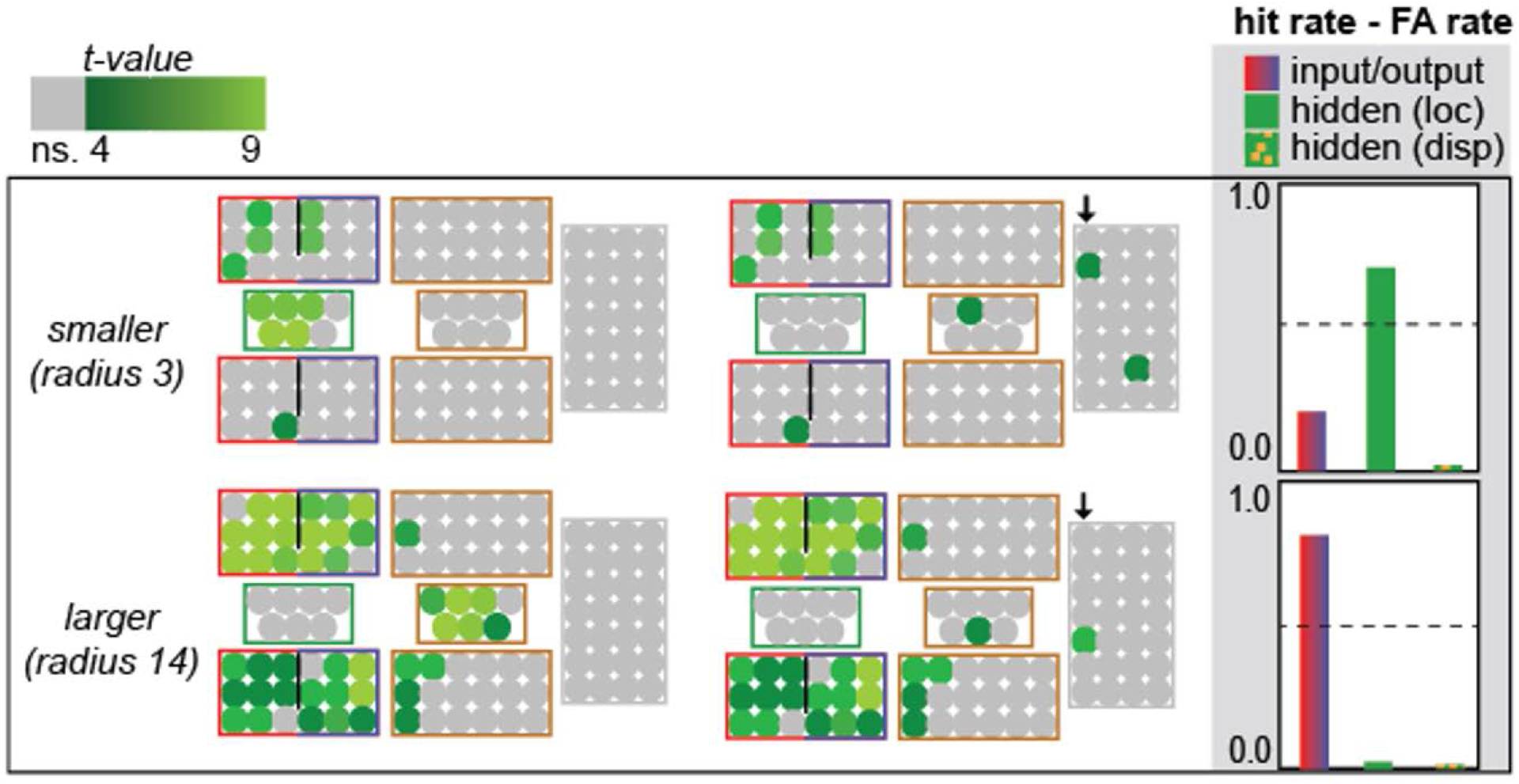
Effect of searchlight size on simulation results.

#### Model results for ridge regularization

Figure 2 in the main paper shows simulation results for L1-regularized logistic regression (LASSO). We here report the results for L2 regularization (ridge regression), which is often the default method for regularizing underconstrained regression problems. Like LASSO, ridge regression returns a vector of coefficients that indicates how the classifier interprets each unit’s activation in generating a predicted class label. To understand which units contribute to the representation of interest and how these units encode information, the coefficients must be interpreted. Interpretation for ridge regression is difficult because the method places a non-zero coefficient on every unit. Intuitively it may seem as though signal-carrying units will attract larger weights, but this is not the case: weights will be small when a unit carries no useful signal, but also when a unit carries useful signal but is highly correlated with many other units. We designated a unit as “selected” by ridge regression if its coefficient was in the top quartile by magnitude across all units. Thus the prior probability of selection was p = 0.25 by definition in each model subject. For each unit, we then assessed whether it was selected more often than expected across model subjects given this base probability, with p < 0.002 uncorrected.

As shown in Figure SI-4, almost no units met this criterion. The same result was also observed for more lax selection criteria (e.g., choosing units in the top third or top half of the magnitude distribution). For this reason we did not show ridge regression results in the main paper figure, and did not consider ridge regression in the fMRI analysis.

**Figure SI-4:**
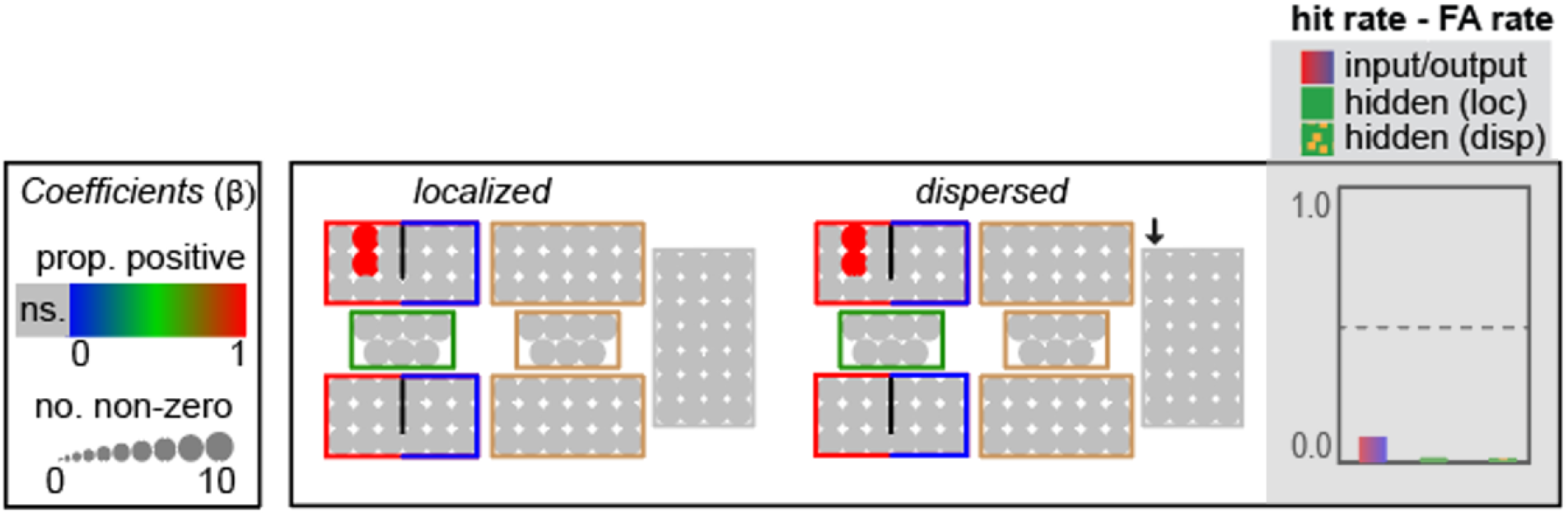
Results of ridge regression applied to simulation data. Colored circles are units that receive a weight in the top quartile more often than expected by chance. Hue indicates the proportion of coefficients that were positive.

This is not to say that ridge regression can never be of use. To the contrary, for scientists who wish to know whether some pre-defined set of voxels contain sufficient information to predict information about a stimulus or task, ridge regression can be very effective. The problem lies, not in determining whether such information is present in a pre-specified set of voxels, but in using the model coefficients to determine which of the voxels are important for the prediction. Thus in the main paper, for instance, we use other statistical methods to first identify voxels “important” for discriminating face from non-face stimuli, then compare the quality of these different solutions by training a ridge-regularized classifier using only the voxels pre-selected by a given method and comparing the hold-out accuracy across methods. This use of ridge regression allows us to employ the same performance measure when comparing the solutions discovered by different methods.

### SI-Imaging

This section reports some additional analyses of the functional imaging data.

#### Effects of searchlight size

Figure 3 in the main paper shows the result of a 9mm searchlight MVPC, which revealed localized signal in the posterior fusiform cortex bilaterally when discriminating faces from other stimuli. This solution differed remarkably from the extended distributed signal discovered by SOS LASSO. The simulations suggest, however, that searchlights can fail to discover noisy signal when they are too small. We therefore conducted the same analysis (discriminating face from non-face stimuli), using a searchlight radius of 15mm, substantially larger than those typically employed in the literature (either 6mm or 9mm). Figure SI-5 shows the result. The larger searchlight revealed more extensive signal extending up into right occipital cortex, but this remained localized within the canonical face system.

**Figure SI-5:**
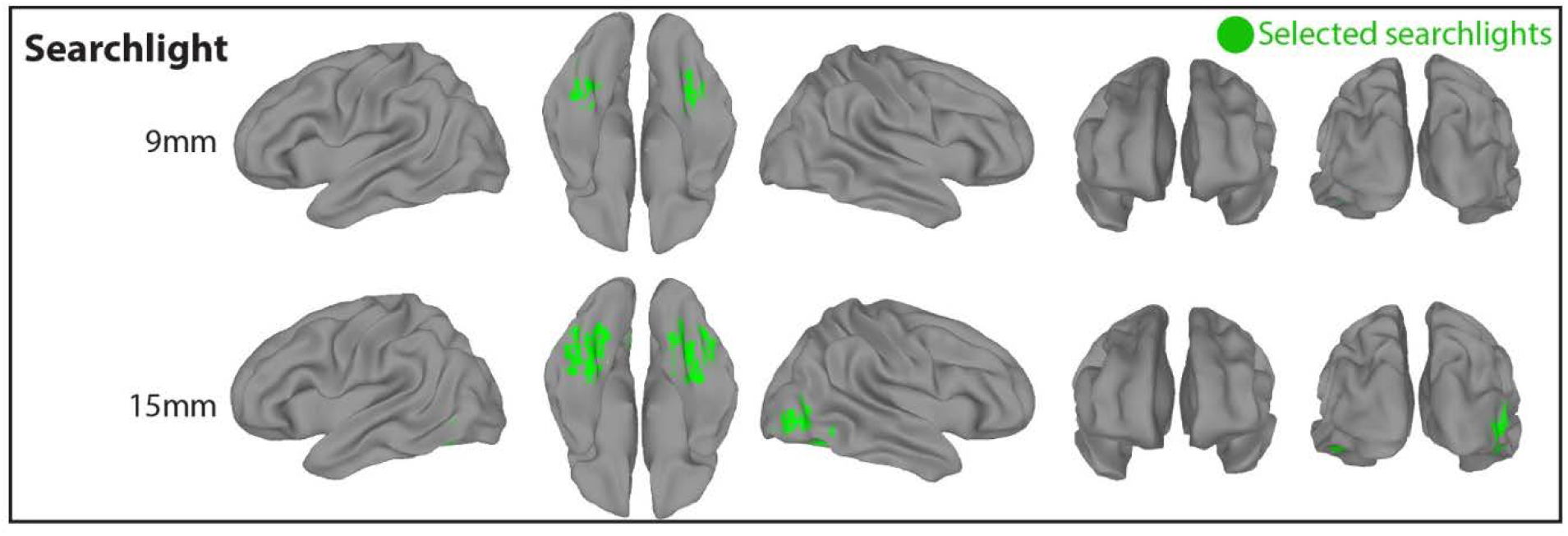
Comparison of results decoding face verus non-face with a standard 9mm searchlight versus a larger 15mm searchlight. The larger searchlight appears to show more distributed signal still confined within the canonical system.

### SI-Denoising: Noise-canceling voxels in whole brain classifiers

An important general concern for whole-brain multivariate decoding arises from the problem of correlated noise across cortex. BOLD estimates across the brain may be simultaneously influenced by some external cause unrelated to the stimulus or task of interest, such as heartbeats, scanner noise, drift in field strength over time, and other sources.^27,28^ These task-unrelated external influences constitute noise in the context of the analysis, but the noise perturbs both signal-carrying and non-signal-carrying voxels in similar ways and so induces partial correlations between these. In this scenario a regression model fit to all voxels together can improve performance by placing weights on both the signal-carrying and non-signal-carrying units. Because the non-signal-carrying voxel activations correlate only with the noise component of the signal-carrying activations, opposing weights on these will “cancel out” the noise perturbing the true signal, effectively boosting the signal-to-noise ratio on the informative voxels. Thus, when noise is correlated across the system, whole-brain classifiers can improve performance by placing non-zero weights on voxels that only encode the correlated noise and are unrelated to the task.

To appreciate this, consider a simple example. Let *x*\* be some true signal, *q*\* be some noise source, *y* = *x*\*, *x*_1_ = *x*\* + *q*\*, and *x*_2_ = *q*\*. Assume that *x*\* and *q*\* are normally distributed with equal mean and variance and are orthogonal. The regression model *y*~*βx*_1_ will explain about half the variance in *y*; the model *y*~*βx*_2_ will explain no variance in *y*; the model *y*~*β*_1_*x*_1_ + *β*_2_*x*_2_ will explain all the variance in *y*, even though *x*_2_ is orthogonal to *y*. This is because the noise *q*\* is shared between *x*_1_ and *x*_2_. The model can use *x*_2_ to cancel the variance associated with *q*\* in *x*_1_, improving the overall fit.

This phenomenon raises the possibility that the highly-distributed signal revealed by SOS LASSO outside canonical face and place systems may arise because the classifier places weights on many units that do not carry signal but are perturbed by noise correlated with that disturbing the true signal encoded within the canonical systems. This concern motivates the two follow-up analyses reported in the main text. First, if extra-system voxels only serve to de-noise the true signal in canonical systems, decoding accuracy should decline when the canonical voxels are removed—yet our decoding accuracy was higher outside the canonical system than within it. Second, if the broad selection of voxels arises because of whole-brain correlated noise, then the same widespread result should obtain regardless of the classification task (since noise in extra-system voxels will also correlated with the noise-component of other signal-carrying voxels). Yet when classifying place versus face/object, SOS LASSO discovers much more localized signal that does not include these regions. Thus, while correlated noise remains a thorny problem for statistical inference generally, in the current case empirical assessments can rule out the possibility that the interesting results arise from this problem.

